# Single-molecule analysis sheds light on cardiac myosin dysfunction due to hypertrophic cardiomyopathy mutation A57D in ventricular myosin light chain-1 (MLC1v)

**DOI:** 10.64898/2026.03.06.710098

**Authors:** Tianbang Wang, Emrulla Spahiu, Matthew Childers, Tim Holler, Kenneth Campbell, Cristobal dos Remedios, Thomas Thum, Theresia Kraft, Michael Regnier, Arnab Nayak, Mamta Amrute-Nayak

## Abstract

Ventricular myosin light chain-1 (MLC1v) is a key structural and function-modulating component of the β-cardiac myosin (βM-II) motor complex. Single-point mutations in MLC1v are linked to severe forms of hypertrophic cardiomyopathy (HCM) and sudden cardiac death (SCD) at a young age. However, the molecular mechanisms underlying the motor dysfunction responsible for HCM phenotype development are not fully understood. Here, we investigated native βM-II motors isolated from septal myectomy sample of an HCM patient, harboring a rare homozygous mutation in MLC1v (A57D). Using a pure population of mutant motors (MUT), and sensitive single-molecule functional analysis approach, we directly assessed the primary functional alterations in βM-II bearing A57D MLC1v mutation. In optical trap single-molecules measurements, the mutant motors displayed increased actomyosin (AM) interaction duration in strongly bound state (t_on_), corresponding to 3-fold reduced AM detachment rate than wild type myosin (WT). The MUT myosin also generated a shorter powerstroke size (δ). Ensemble average analysis of AM interaction events demonstrated that both the first powerstroke (δ_1_) associated with Pi release and the second powerstroke (δ_2_) linked to ADP release were reduced in MUT myosin. Moreover, the increased actomyosin cross-bridge stiffness in the AM.ADP state was observed for MUT compared to WT motors. Consistent with slower AM detachment rate and shorter stroke size, reconstituted human mutant βM-II displayed slower actin filament gliding speed. Alterations in sarcomere-level mechanics included increased Ca^2+^ sensitivity of force generation and prolonged relaxation time, as predicted by FiberSim modelling. Molecular dynamics simulations indicated that the substitution of alanine by aspartate altered MLC1v interactions with myosin heavy chain (MyHC) and light chain 2 (MLC2v), affecting the curvature and flexibility of the lever arm. Overall, these studies establish the molecular mechanism underlying the primary myosin dysfunction due to A57D MLC1v mutation and further highlight the crucial role of MLC1v-mediated regulation of myosin function. Understanding the precise changes in the mutant myosin’s biomechanical properties is directly relevant to comprehending the initial triggers for pathological cardiac remodeling in HCM patients and designing tailored therapeutic interventions.

## Introduction

Hypertrophic cardiomyopathy (HCM) is an autosomal dominant disorder with an estimated prevalence of 1 in 200 individuals worldwide ^1, 2^. The heart disease is characterized by thickening of the left ventricular wall (LV) and the inter-ventricular septum (IVS), increased interstitial fibrosis and cardiomyocyte disarray. Thickened LV and IVS result in small chamber size and reduce the end-diastolic blood volume (EDV). The hypertrophied heart exhibits preserved or increased ejection fraction (EF) and suffers from outflow tract obstruction in about two-thirds of HCM cases and impaired diastolic function in almost all cases. HCM is often linked to sudden cardiac death in young individuals and competitive athletes ^3^. Given the high prevalence and mortality linked to HCM, there is an urgent need for better understanding of the primary molecular level dysfunction to find effective therapeutic strategies to treat HCM patients.

HCM is primarily caused by inherited mutations in sarcomeric proteins, including the components of β-cardiac myosin (βM-II), which is a hexameric holoenzyme comprising of two myosin heavy chains (MyHCs) and four light chains (LCs). Each MyHC contains three main structural domains: 1) a catalytic domain containing ATP and actin binding sites, 2) a lever arm or light chain binding domain, where essential (ELC/MLC1v) and regulatory light chains (RLC/MLC2v) associate with the IQ motifs of the heavy chain, and 3) a coiled-coil tail domain, which forms bipolar myosin thick filaments. As the main force-generating motor expressed in the human heart ventricle, βM-II drives the cardiac contraction through the ATP-dependent interaction with actin (Figure 1 Scheme). During the cross-bridge cycle, the nucleotide-dependent transition from weak to strong binding states of the actomyosin (AM) complex generates the power stroke and movement. The light chains stabilize the ∼ 9 nm long alpha-helical structure of the lever arm. Apart from their structural role, the light chains are also implicated in regulating the AM cross-bridge cycle and influencing the Ca^2+^ sensitivity of force generation in an isoform-dependent manner ^4^, through phosphorylation of RLC and the interaction of the N-terminus of ELC with actin filaments ^5–7^. The importance of ELC in myosin function are highlighted by studies showing that their removal from myosin motors reduces force and causes loss of actin filament movement ^8^. Several HCM-causing missense mutations have been found in MLC1v ^9, 10, 11^. Although rare, i.e., < 5 %, MLC1v mutations are associated with malignant forms of HCM and sudden cardiac deaths (SCD) at a young age ^12^. These findings further reinforced the importance of MLC1v as a crucial modulator of myosin function. The underlying mechanistic details of how a specific MLC1v mutation mediated βM-II malfunction leads to disease phenotype remains unresolved.

**Figure 1.**
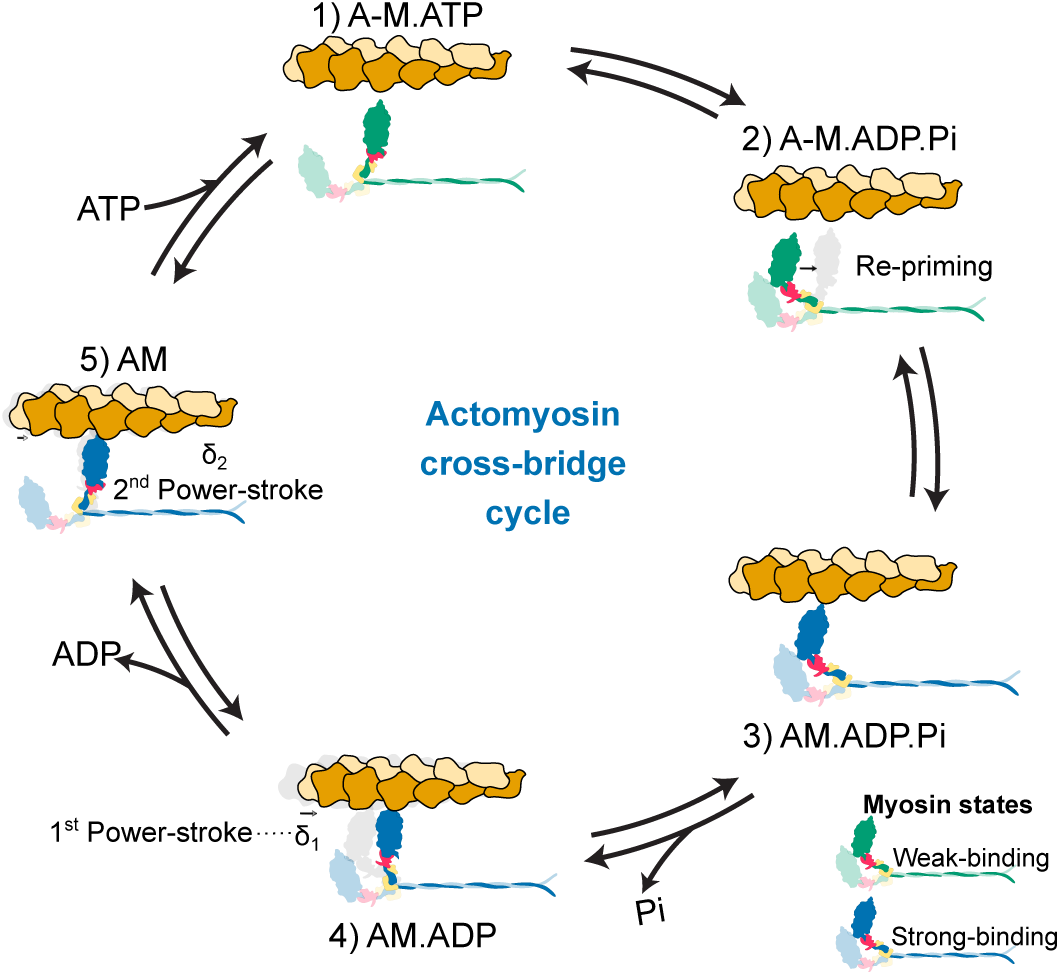
The simplified schematic summarizes the essential conformational states during actomyosin crossbridge cycling. Strongly- and weakly-bound AM states are indicated with blue and green- myosin heads.

Human myocardial tissues from HCM patients are valuable sources for scrutinizing inter-and intracellular structural remodeling, as well as altered organelle and biomolecular functions, helping to understand the overall HCM pathomechanism. However, studying the functional effects of HCM-related mutations in βM-II motor components using patient tissue is hampered by several factors: 1) limited availability of myectomy or explanted heart tissue, and 2) patients being heterozygous for HCM mutations; thus, both mutant (MUT) and wild-type (WT) myosin motors are incorporated into the contractile apparatus of the ventricular myocardium. The expression of mutant protein ranges between 10 to 75% of the total motor protein ^13^. The co-expression of MUT and WT motor proteins limits the ability to identify the direct functional effects of mutations in ensemble average measurements of myosin, and subtle functional changes can be readily missed. In case of low abundance of the mutant protein, even significant functional effects become undetectable in ensemble measurements.

In previous studies, various alternative strategies were employed to understand the HCM pathology. Transgenic animal models, including mouse and rabbit, were generated to study various HCM-associated mutations in MyHC and LCs, allowing characterization of the resulting HCM phenotype ^14–17^. To explore mutation-specific motor characteristics, human MLC1v were reconstituted with bovine cardiac myosin II ^18^, and patient-derived slow-twitch muscle fibers were employed to evaluate the ATPase activity, force generation, and fiber stiffness for the corresponding HCM mutations ^19, 20^.

Although these approaches have substantially advanced our understanding of HCM pathogenesis, a major limitation persists: the myosin complexes analyzed often contain subcomponents from mixed species and varying isoform compositions. For example, in transgenic animal models, MUT human MLC1v is assembled with mouse α-MyHC and MLC2 ^17^. Notably, the atrial myosin comprising α-MyHC isoform exhibits higher ATPase activity and distinct mechanical features than the ventricular myosin with β-MyHC isoform. Moreover, slow-twitch muscle fibers, often considered a substitute for cardiomyocytes, express different isoforms of troponin (slow skeletal ssTnT and TnnI) and myosin binding protein-c in humans. Consequently, the specific molecular composition and isoform identity of myosin subunits, as well as other sarcomeric proteins in cardiomyocytes, are expected to substantially influence the measured contractile properties.

In these circumstances, studies on human cardiac tissue-derived ventricular motors are key to understanding the normal or altered function of the cardiac cells under pathological conditions. Here, we employed the rarely found homozygous mutation in MLC1v (A57D) to investigate the primary dysfunction of the βM-II motors that led to the HCM phenotype in the patient. The point mutation A57D is located at the α-helix of the first EF-hand Ca^2+^ binding motif of MLC1v, where alanine is an evolutionarily conserved residue. The American College of Medical Genetics and Genomics (ACMG) categorized the pathogenicity of the MYL3 c.170C>A as a variant of uncertain significance (VUS). As homozygous HCM mutations are rare, access to this biopsy specimen provided us an opportunity to study the native pure population of mutant motors. We observed alterations in biomechanical properties at the individual molecule level that included reduced actomyosin detachment rate, powerstroke size, and increased stiffness of the MUT myosin. Molecular dynamics simulations revealed structural changes in the MLC1v and its interactions with MyHC and MLC2v as a consequence of the mutation, leading to net biomechanical changes in the MUT motor. Furthermore, we probed the reliability of the reconstituted human βM-II by directly comparing the native *vs.* reconstituted wild-type and mutant myosins. The functional resemblance indicates that the reconstituted myosin may serve as an alternative source for studying other light chain mutants in the future where patient-derived samples are not available.

Overall, this study elucidates the underlying mechanism of the A57D MLC1v mutation-induced HCM pathology. It provides a rationale for the preferred motor protein source for subsequent investigations, recreating myosin function/dysfunction in the human heart. The benefits of the reconstitution approach extend to the examination of other missense mutations in genes encoding sarcomeric proteins to unravel phenotypic variations and thereby design targeted therapeutic strategies.

## Results

We employed the excised tissue from septal myectomy of a 52 year old male HCM patient, harbouring a homozygous mutation in the *MYL3* gene (c.170C>A) that encodes ventricular myosin light chain 1 (MLC1v), to examine the disease phenotype and assess the impact of the MLC1v point mutation (A57D) on β-cardiac myosin (βM-II) function. An age and sex-matched left ventricular tissue sample from a 56-year-old male donor (non-diseased heart) was selected as a control wild-type (WT) subject for comparison with the patient tissue. The small sized patient myectomy sample required careful consideration; therefore, we adopted the workflow schematized in Figure S1 to optimize the usage of the human tissue sample for various experiments.

### Tissue level cardiac remodelling in patient myocardium

To probe the extent of cardiac remodeling at the cellular level, we performed immunohistochemistry on the cryosections from the donor’s and the patient’s cardiac samples. The tissue sections were stained with different antibodies, including Z-disc marker α-actinin, gap junction markers connexin-43 and N-Cadherin, and the connective tissue marker collagen-I, to visualize the sarcomere and myofibrillar organization, cellular morphology, compaction, cellular alignment, and fibrotic tissue in interstitial spaces. A distinct diseased phenotype was observed at the cardiomyocyte level as shown in Figure 2. While well-aligned, compact cardiomyocytes were evident in the donor probes, the patient’s cardiac cryoslices showed cardiomyocyte disarray and distinct larger fibrotic tissue areas (Figure 2A and B). Moreover, the gap junction protein connexin-43 that marks the cell-cell junctions, and cell adhesion molecule N-Cadherin showed altered distribution and proteins in puncta form in diseased tissue in contrast to homogenous distribution at cell-cell connection surfaces in donor tissue slices (Figure 2C &D). In the donor sample, regularly spaced, distinct, homogenous localized adhesion molecules resided between the cell-cell contact points. The redistribution of connexin-43 and N-Cadherins, which are important for cell-cell communication through excitation-contraction coupling, indicated a possible loss of coordination between cells, which is essential for the synchronous beating of the heart.

**Figure 2.**
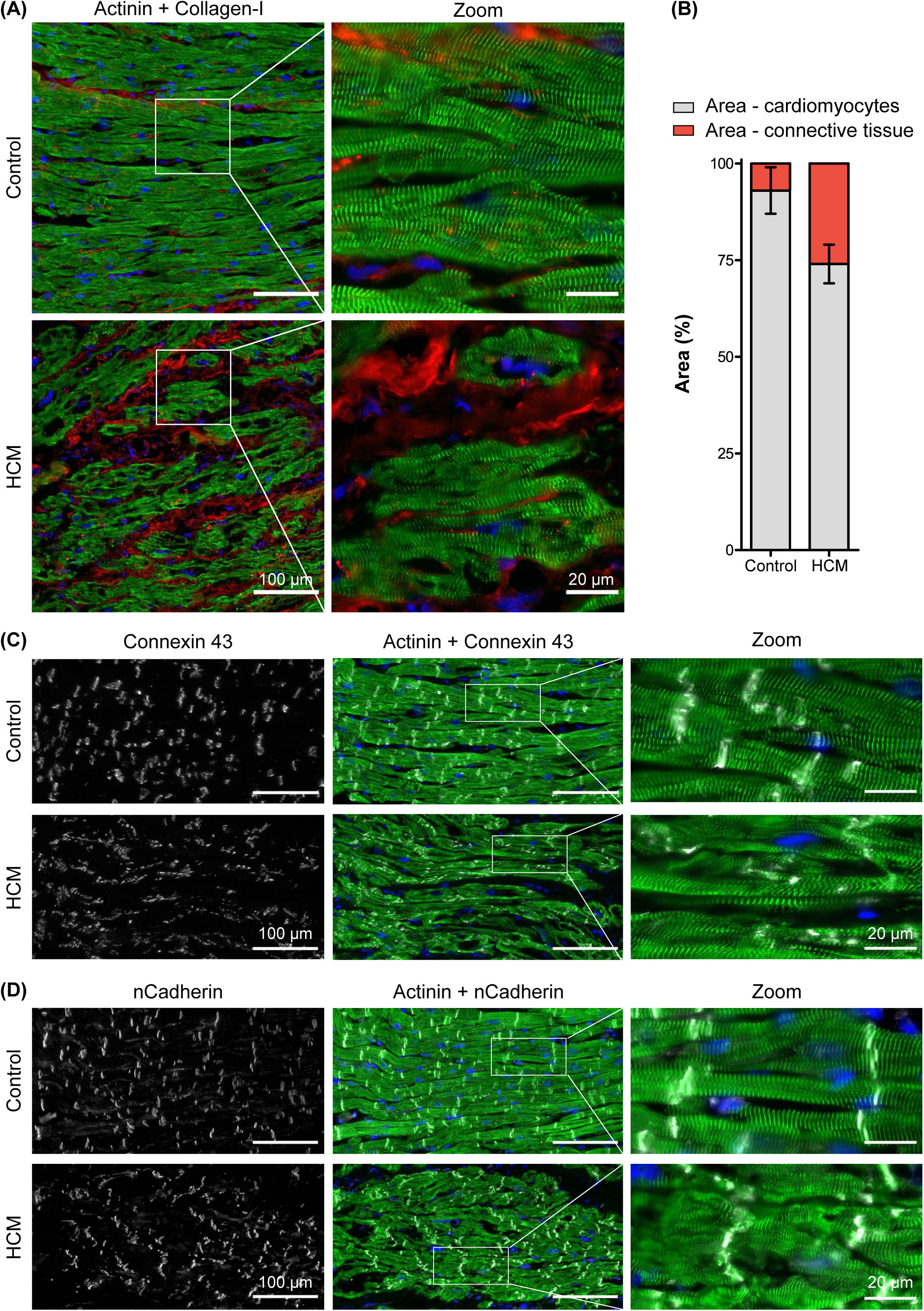
Tissue level cardiac remodeling in patient myocardium. **A)** Donor and patient cardiac tissue sections were stained for α-actinin (green) to probe the cardiomyocytes Z-disk organization, collagen-I (red) to visualize fibrosis, and with DAPI (blue) to stain for nuclei. Images are merged to combine the different antibodies and DAPI staining, i.e., cardiomyocytes (green) and fibrotic area (red). White boxes in all figures indicate the zoomed-in view to show the cardiomyocytes and fibrotic regions. **B)** Analysis of percentage of fibrotic area in the tissue slices. ImageJ tool was used to quantify the areas covered by cardiomyocytes and fibrosis. **C-D)** Cardiac tissue slices were probed for gap junction proteins, connexin-43 or n-cadherin (white) and α-actinin (green) for cardiomyocytes in donor and patient tissue slices as indicated. Scale bars are indicated in respective images.

### Heterogeneity at the tissue level

A comparison of healthy and diseased samples revealed changes in the myosin heavy chain isoform composition, i.e., increased α-MyHC expressing cells in the patient’s tissue sections (Figure 3A and B). While the majority of cells appeared exclusively α- or β-MyHC positive, in some cardiomyocytes, both α-MyHC and β-MyHC were expressed. Therefore, we further checked whether the α- and β-MyHC present in the same cardiomyocytes are even assembled in the same sarcomeres (Figure 3C and D). We observed the co-existence of α- and β-MyHCs both in donor and patient tissue sections, albeit at low levels in donors than in patient tissue. The line profiles for α- and β-MyHC staining further confirmed the intra-sarcomeric presence of both the heavy chains. The intensity for the gap junction protein n-cadherin can also be seen as misplaced at several sites along the cardiomyocyte in patient tissue (Figure 3D), further indicating the cardiomyocyte disarray. Collectively, the presence of both α-MyHC and β-MyHC was confirmed, suggesting a heterogeneous mixture of motors within the sarcomeres.

**Figure 3.**
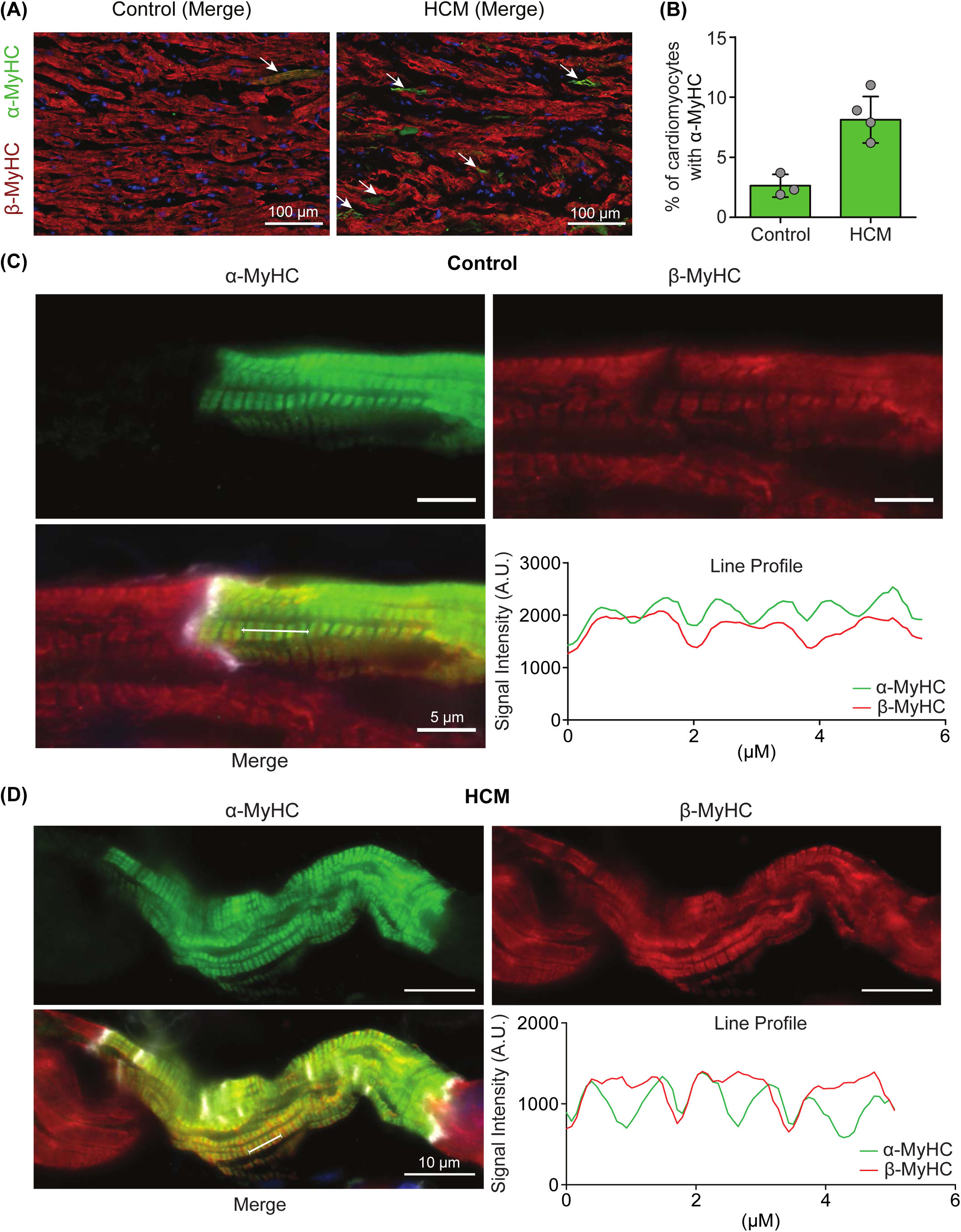
Heterogeneity at the tissue level. A) Immunostaining of donor and patient-derived tissue sections with α-MyHC (green) and β-MyHC (red) specific antibodies. White arrows indicate α-MyHC positive cells. Only one cardiomyocyte can be observed with distinct α-MyHC expression (arrow) in donor tissue *vs.* multiple cells in patient. **B)** The quantification shows the increased number of cardiomyocytes with α-MyHC signal in patient cryosections. Each dot represent a cryosection. **C-D)** Individual cardiomyocytes displaying the co-expression of α-MyHC (green) and β-MyHC (red). N-cadherin staining (white), marks the connections between neighboring cardiomyocytes. The overlapping red and green signal within the sarcomeres are further shown by the signal line profile plot in the right panels for both donor and patient tissues. About 2 µm sarcomere length is evident. A peak with wider intensity distribution are observed with β-MyHC (red line), and two peaks within 2 µm distance with α-MyHC (green line). Note that the epitope for the antibody binding site is in the tail region in β-MyHC and close to the motor domain in α-MyHC, thereby, resulting in observed characteristic intensity distribution.

### Composition of cardiac myosins from donor and HCM patient

In the next step, wild-type and mutant βM-II molecules were extracted from donor and HCM patients’ IVS tissue samples, respectively (Schematic 4A). For control, atrial myosin αM-II was isolated from the human atrial appendage. The extracted myosin was assessed for purity and the presence of different isoforms of myosin heavy and light chains (Figure 4). In adult humans, IVS tissue predominantly expresses β-cardiac myosin heavy chain (β-MyHC). As shown in Figure 4B, no α-MyHC (atrium-specific myosin heavy chain isoform) was detectable in the Sypro-Ruby-stained gel, which was employed to inspect the heavy chain isoforms for donor (WT) and patient tissue-derived myosin (MUT). In the WT atrial tissue sample, however, predominantly α-MyHC and a small fraction of β-MyHC were visible. The light chain (LC) composition was also examined, as shown in Figure 4C. As expected, ventricle-specific MLC1v and MLC2v were observed in IVS tissue of the HCM patient-derived myosin (Figure 4C). Strikingly, apart from MLC1v, a faint band corresponding to MLC1a, that is an atrial myosin light chain-1, was also observed. Note that in the immunofluorescence assay, we observed atrial MyHC in the cryosections of patient-derived tissue (Figure 3A). The α-MyHC, however, was not detectable in the mutant sample in the gel image. One possible reason may be that the intensities corresponding to the α-MyHC band remained below the detection limit for the amount that was loaded onto the gel.

**Figure 4.**
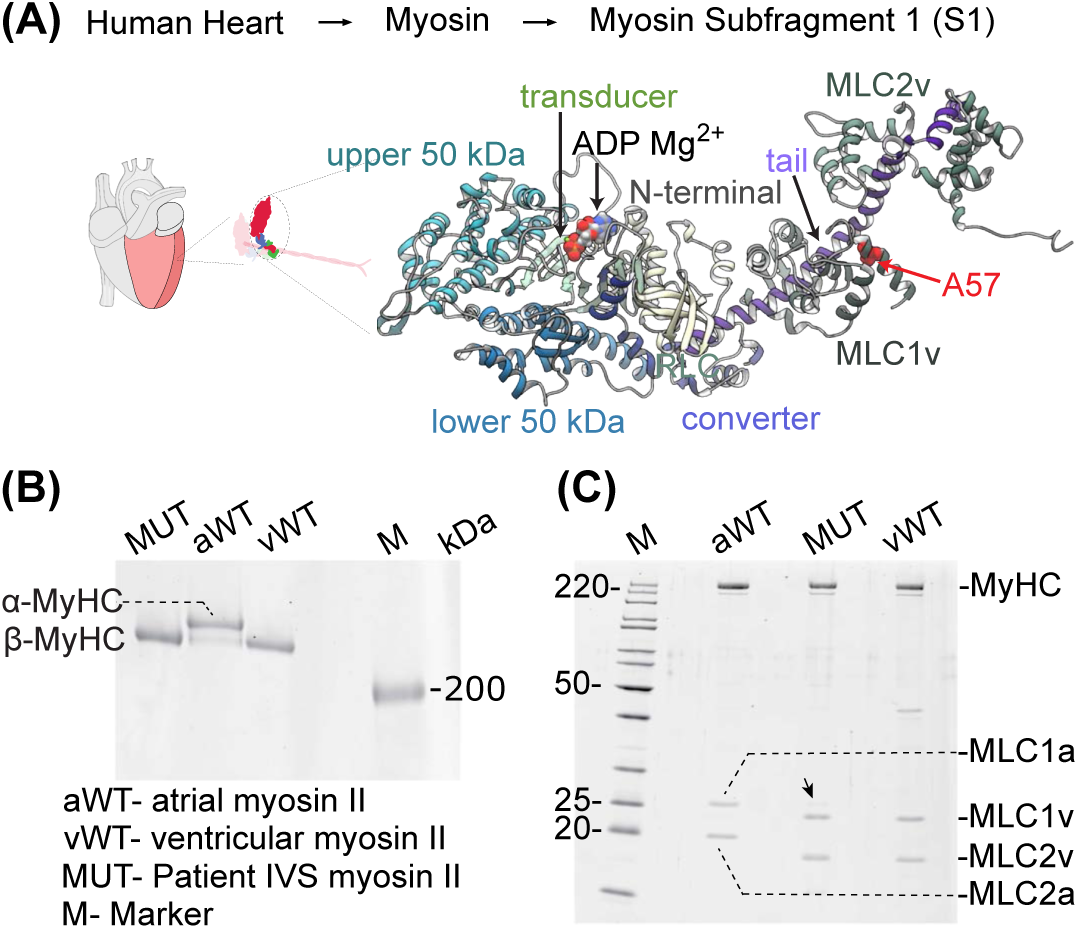
Heavy chain and light chain composition of the cardiac myosins obtained from patient and donor cardiac samples. **A)** Illustration of heart and dimeric cardiac myosin, together with human βM-II subfragment 1 structure indicating the A57D MLC1v mutation site. The myosin motor domain is organized into four domains (N-terminal domain, upper and lower 50 kDa domains, and the converter domain) that are arranged around the nucleotide binding pocket and central β-sheet (that contains the transducer) are shown. The essential (MLC1v) and regulatory light chains (MLC2v) bind to ‘IQ’ motifs within the myosin lever arm region. **B)** MyHC isoform gel showing donor and patient ventricular samples having only β-MyHC isoform, while atrial sample having ∼20% β-MyHC (estimated from the densitometry analysis). **C)** Gel showing chamber-specific light chains from atrium and ventricle of donor samples. In MUT sample, the black arrow indicates the presence of a light intensity band of corresponding molecular weight as atrial MLC1a. Abbreviations: α-MyHC and β-MyHC - atrial-specific alpha or ventricle-specific β myosin heavy chains around 220 kDa. MLC1v and MLC2v - ventricle myosin specific light chain-1 (LC1) or essential light chain (ELC) and light chain-2 (LC2) or regulatory light chain (RLC), respectively. MLC1a and MLC2a are atrial myosin-specific LC1 and LC2, respectively.

### Single-molecule function analysis of WT and MUT myosins

To understand the primary motor dysfunction caused by the mutant MLC1v, motor properties were investigated using single-molecule optical trapping analysis (Schematic Figure 5A). The motor function determining parameters include ‘lifetime’ of actomyosin interaction and ‘displacement’ of the actin filament during the ATP-dependent actomyosin crossbridge cycle. The duration of strongly-bound actomyosin states ‘t_on_’, the stroke size (δ), and actomyosin crossbridge ‘stiffness’ are contributing factors to the sarcomere shortening velocity and governs the force-generating ability of the myosins. Using optical trapping, we directly measured the kinetics of actomyosin interaction and the nanoscale mechanical parameters, i.e., the stroke size and stiffness of individual motor molecules, with high spatiotemporal resolution.

**Figure 5.**
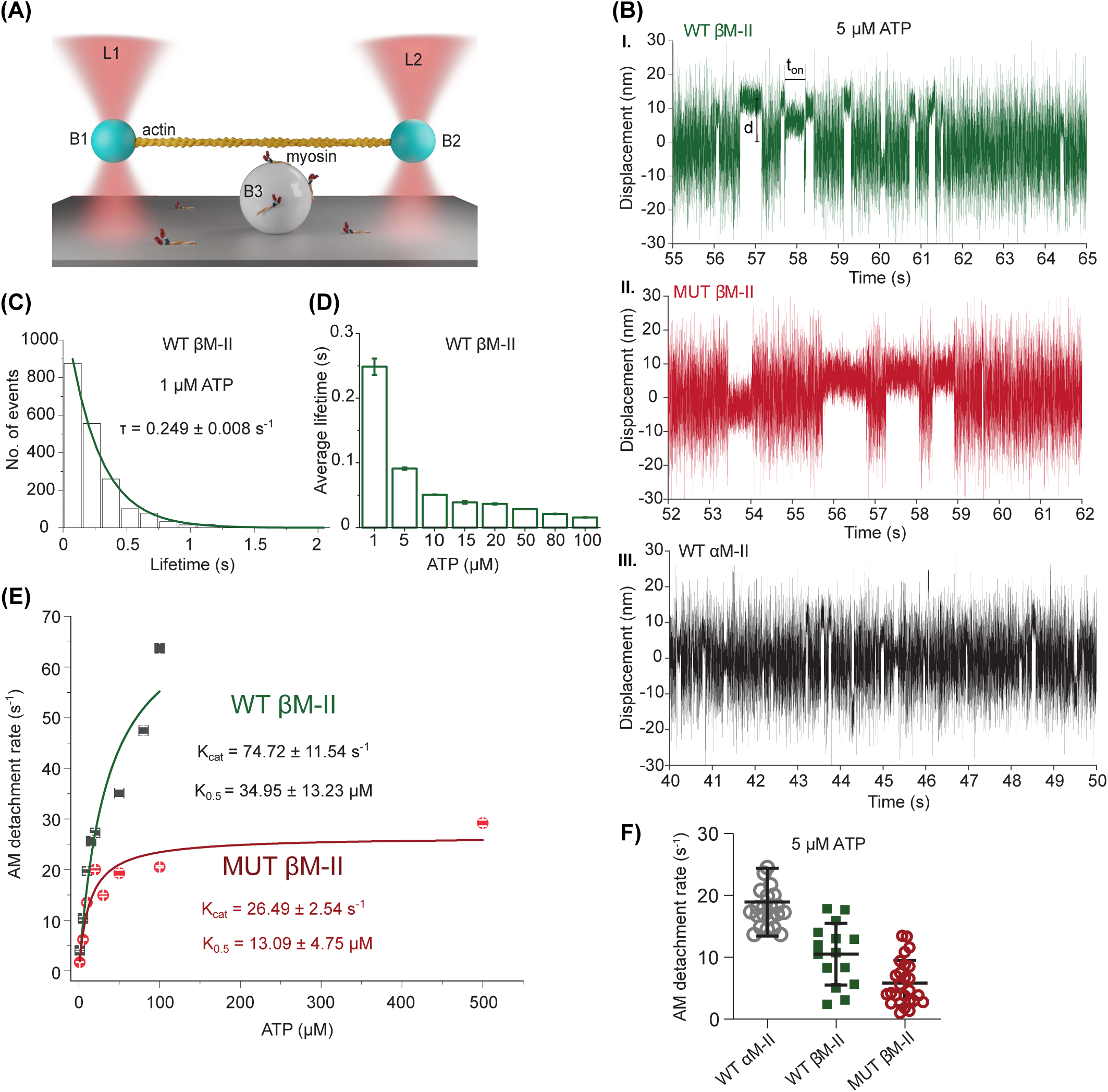
Slower AM detachment kinetics for MUT myosin. **A)** Optical trap setup for three-bead assay. Note that the various components of the setup are not drawn to scale. Actin filaments are suspended between the two optically trapped beads (bead 1 and bead 2). The motor is adsorbed on the glass bead (bead 3) immobilized on the surface. The two bead positions are registered using a quadrant detector to detect the interaction between actin and myosin. **B)** The original data traces displaying the bead position signal over time. **I-III)** Representative original data traces for WT βM-II, MUTβM-II and αM-II showing single myosin molecules interacting with an actin filament at 5 µM ATP. Reduction in the amplitude of bead displacement or Brownian motion, indicates AM interaction events. t_on_ - the duration of AM association. The vertical green line segment illustrates the displacement (d) from the average unbound (0) to the average bound position. **C)** The histogram comprising single molecule AM interaction lifetimes at 1 µM ATP. More than 1000 individual AM events were plotted to estimate the average lifetime in sec (τ). **D)** Average AM interaction event lifetime (τ) at increasing [ATP] are plotted. **E)** AM detachment rate (1/τ) for the corresponding ATP concentrations shown in **(D)** were estimated. ATP concentration dependence of AM detachment rate is plotted and fitted to a Michaelis Menten’s equation to determine the maximum detachment rate (*k_cat_*) and the ATP concentration at half-maximal detachment rate (K_0.5_). **F)** Comparison of the AM detachment rate from individual motor molecules for atrial myosin, WT αM-II, WT βM-II, and MUT βM-II. N-Number of molecules, n – no. of events. For WT myosin, at 1 µM ATP, N = 13, n = 1791, at 5 µM ATP N=15 and n= 2363, at 10 µM ATP N= 25, n= 5042, at 15 µM ATP N= 13, n= 2081, at 20 µM ATP N= 8, n= 1021, at 50 µM ATP N= 15, n= 1022, at 80 µM ATP N= 13, n= 1191, at 100 µM ATP N= 9, n= 748. For MUT myosin, at 1 µM ATP, N = 23, n = 1223, at 5 µM ATP, N= 28 and n= 2582, at 10 µM ATP N= 24, n= 3995, at 20 µM ATP N= 11, n= 1717, at 50 µM ATP N= 17, n= 1521, at 100 µM ATP N= 31, n= 4575, and at 500 µM ATP N= 33, n= 4159.

### A57D mutation altered the actomyosin detachment kinetics

We analyzed individual cardiac myosin molecules for their interaction with actin as described in earlier reports ^21, 22^ (examples traces shown in Figure 5B). In displacement over time traces obtained from WT βM-II, MUT βM-II, and WT αM-II (Figure 5B), the differences in the durations of actomyosin binding at the same ATP concentrations (5 µM) were evident. The duration of the AM interaction comprises the lifetime of the ‘ADP-bound’ state and the nucleotide-free ‘rigor’ state. Following the transition of myosin from the weak to the strong actin-bound state, rapid Pi release and powerstroke generation occur. We assume that the lifetime of Pi-bound AM state (AM.ADP.Pi) is too short to contribute substantially to the observed duration of AM interaction in trap experiments with βM-II. Note that we are unable to resolve the early step of weak to strong transitions, i.e., initial AM binding and generation of powerstroke, which is reported to occur in <2 ms ^23^.

The event lifetimes for WT βM-II were measured at various ATP concentrations, as shown in Figure 5C and D. The average AM bound lifetime (τ) decreased with increasing ATP concentrations in the measured range of 1 µM to 100 µM, as presented in Figure 5D. The AM detachment rates are derived from the average AM association duration (1/τ). At high [ATP], the contribution from the lifetime of the rigor state becomes insignificant, and the AM detachment is primarily limited by the ADP release or binding of the ATP to myosin, resulting in the actomyosin dissociation. Increasing ATP concentration shortened the duration of the rigor state and thereby increased the rate of AM detachment (Figures 5E). Note that for WT βM-II, 100 µM ATP was the upper limit, as the interaction events were too short to be detected with our spatial resolution when higher [ATP] were used. For the MUT βM-II, however, from 50 µM ATP, the detachment rates remain essentially unchanged, and therefore, the measurements at even 500 µM ATP were possible. The [ATP] dependence of the AM detachment rate (Figure 5E) revealed the maximal detachment rate (k_cat_) of 74.5 s^-1^, and the ATP concentration to obtain half-maximal detachment rate, K_0.5_ of 34.9 µM for WT myosin. A significant difference was observed for the MUT βM-II. Beyond 20 µM ATP, the detachment rate did not increase markedly as observed for the WT βM-II. For the MUT βM-II, k_cat_ of 26.5 s^-1^ and K_0.5_ of 13 µM was estimated. These results indicated a ∼3-fold slower detachment of myosin from actin and the improved sensitivity of myosin for ATP by about 2.5-fold (as indicated by the lower K_0.5_ for mutant motors). The linear regression of the detachment rates over the ATP concentration range from 1 to 10 µM ATP yielded the second-order rate constant of ATP binding of 1.75 ± 0.08 µM^-1^s^-1^ and 1.31 ± 0.10 µM^-1^s^-1^ for WT and MUT βM-II, respectively. Comparable rate constants of ATP binding for both WT and MUT βM-II suggest that the differences in the maximal detachment rate represent the altered rate of ADP release due to the A57D mutation in MLC1v.

We employed atrial-specific α-cardiac myosin (αM-II) as a control, since small amounts of α-MyHC were found in the ventricular tissue sections (Cf. Figure 3A). αM-II is known to display 2-3-fold faster velocity than ventricular myosin ^24, 25^. A clear difference was observed between the actomyosin detachment rates for α- and WT βM-II, with a 2-fold higher rate of about 19 s^-1^ for αM-II and 10 s^-1^ for βM-II, at 5 µM ATP. The difference between the αM-II and MUT βM-II was about 4-fold at 5 µM ATP (Figure 5F). These results indicated that the majority of the analyzed molecules contained βM-II in mutant myosin preparation.

### A57D mutation reduced the myosin’s powerstroke size

Next, we determined the powerstroke size of the human cardiac myosins. Individual binding events were analysed for distance from the mean bead displacement position (cf. data trace in Figure 5B) and plotted to evaluate the stroke size. The shift-of-histogram method ^26^ was used to determine the average stroke size or the displacement of the actin filament by the myosin molecule during individual AM interactions. As shown in Figure 6A and B, the MUT myosin showed a significantly shorter stroke size of 3.4 nm as compared to the WT myosin with a stroke size of about 5 nm. WT αM-II exhibited a stroke size of about 4.23 nm under identical [ATP] (Figure 6C).

**Figure 6:**
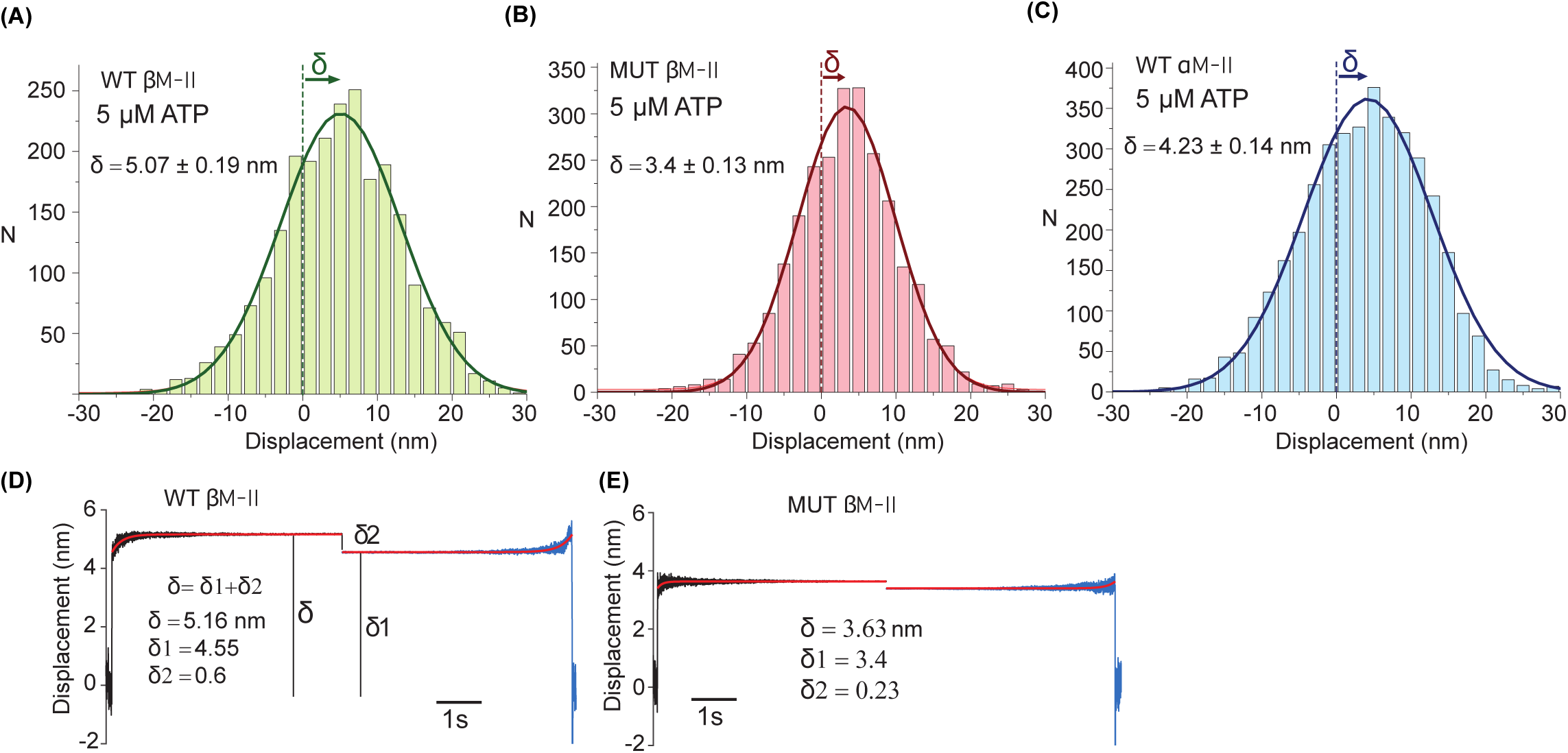
Powerstroke size of WT vs MUT. β**-cardiac myosin.** Stroke size estimated using two methods, namely ‘histogram-shift’ and ‘ensemble-average’ method. **A-C)** The displacements from individual actomyosin interaction events plotted in a histogram is fitted with a Gaussian function to derive the average stroke size. The dotted line indicates the average dumbbell position (0) in the unbound state and the horizontal arrows show the average displacement of 5.07± 0.19 nm and 3.4 ± 0.13 nm for WT and MUT β-cardiac myosin as it interacts with actin filament. For αM-II, average stroke size was 4.23 ± 0.14 nm. **D and E)** AM interactions were synchronized at the beginning and end of the events. Ensemble averaging used to estimate the first and second powerstroke size, δ_1_ and δ_2_, for WT vs MUT β-cardiac myosin. For WT βM-II, N=15 and n= 2363, Mut βM-II N= 28 and n= 2582.

The total stroke size is expected to include substeps associated with the release of Pi and ADP from the active site. The 1^st^ powerstroke, linked to the Pi release, is the beginning of the mechanical interaction between actin and myosin. Within the AM-bound period, the 2^nd^ powerstroke associated with the ADP release occurs as myosin transitions from AM.ADP to AM state (c.f. ATPase Scheme Figure 1). The amplitude of the Brownian motion of the trapped bead is larger than the stroke size. Given a rather small overall stroke size of ∼5 nm, it is difficult to extract the substeps from normal displacement-time records. Moreover, as the first powerstroke occurs in less than 2 ms after binding to the actin filament, the damping effect of the trapped bead masks the myosin’s conformational change following the initial AM attachment. To resolve the substeps, we adapted the ensemble-averaging method^27^, in which sufficiently long events at the start and end of the events were synchronized to reveal hidden information in the noise with high precision. We employed a Matlab-based program, SPASM ^28^, to synchronize the events as shown in Figure 6D-E. At lower ATP concentrations of 5 µM, the bound durations of AM are expected to contain sufficiently long time traces to derive the amplitude of 2nd stroke (i.e., lever arm rotation corresponding to transition from AM.ADP to AM state). For WT βM-II, the analysis revealed the total stroke size of 5.16 nm and amplitude of each step to be 4.55 nm and 0.6 nm for the 1^st^ and the 2^nd^ stroke, respectively. For MUT, total stroke size of 3.63 nm and amplitude of each step of 3.4 nm and 0.23 nm for the 1^st^ and the 2^nd^ stroke, respectively was estimated. Thus, the MUT MLC1v affected the overall working stroke of MUT βM-II by shortening the amplitude of both the 1^st^ and 2^nd^ substeps.

### A57D mutation altered the AM crossbridge stiffness in the ADP bound state

The stiffness of the motor molecules directly influences their force generation capacity. As a key mechanical parameter, we examined if the actomyosin cross-bridge stiffness is affected by the A57D mutation in MLC1v (Figure 7). The details of stiffness measurement are provided in the M &M section. We previously demonstrated that the AM crossbridge stiffness in the ‘ADP-bound’ state is lower as compared to the AM ‘rigor’ state ^21^. The two states were acquired by adjusting the ATP concentrations such that the measured AM bound duration comprised primarily ‘AM-ADP’ at high [ATP] or ‘AM rigor’ state at low [ATP].

**Figure 7.**
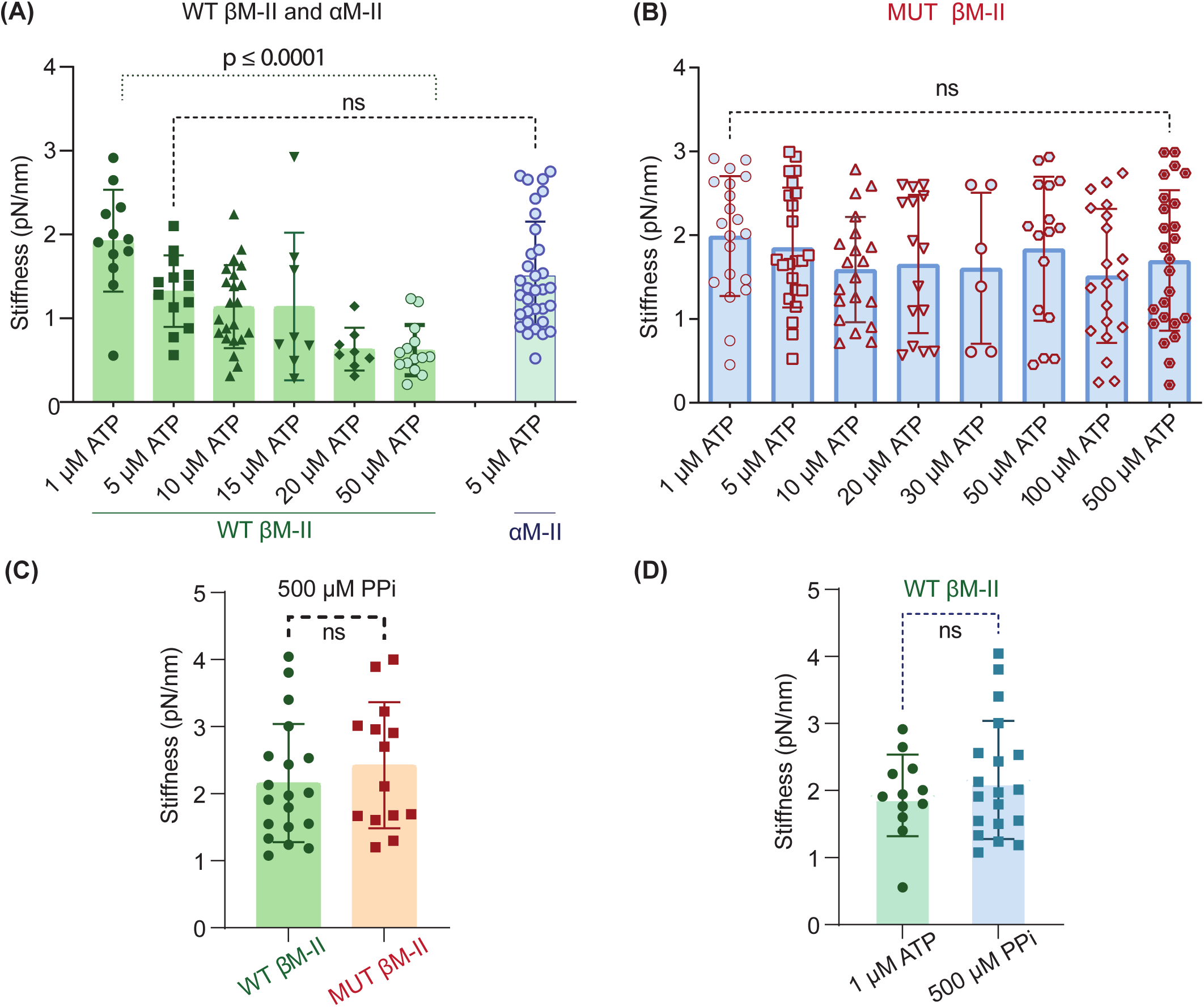
The rigidity of the AM crossbridges was estimated using the variance-covariance method. **A)** The scattered data plots show stiffness measured at various ATP concentrations for WT β- and α-cardiac myosin, and **B)** MUT β-cardiac myosin. Each data point represents the stiffness of an individual myosin molecule as it interacts with the actin filament. **C)** AM crossbridge stiffness in the presence of 500 µM PPi for WT and MUT βM-II (P=0.41). **D)** WTβM-II - comparison between 1 µM ATP and 500 µM PPi (P=0.43). Error bars are standard deviations (SD) from the mean values. Ns-non significant. PL0.0001 for the stiffnesses compared between 1 µM and 50 µM ATP condition for the WT β-cardiac myosin. Statistical analysis using unpaired t-test. For WTβM-II, at 1 µM ATP, N = 12, n = 1021, at 5 µM ATP N=14 and n= 1754, at 10 µM ATP N= 22, n= 3454, at 15 µM ATP N= 8, n= 1369, at 20 µM ATP N= 8, n= 901, at 50 µM ATP N= 14, n= 528. For MUT βM-II, at 1 µM ATP, N = 20, n = 1336, at 5 µM ATP N=23 and n= 2228, at 10 µM ATP N= 20, n= 2599, at 20 µM ATP N= 9, n= 984, at 30 µM ATP N= 6, n= 379, at 50 µM ATP N= 16, n= 1182, at 100 µM ATP N= 20, n= 2721, at 500 µM ATP N= 25, n= 1648.

Consistent with our earlier studies on rabbit cardiac myosins ^7^, ATP concentration dependence of the stiffness was observed for human WT βM-II (Figure 7A). Lower stiffness of 0.62 pN/nm was estimated at 50 µM ATP and ∼3 fold higher stiffness of 1.93 pN/nm at 1 µM ATP. For αM-II at 5 µM ATP, we estimated the average stiffness of the AM crossbridge state to be 1.33 pN/nm. Thus, stiffness of WT βM-II was similar to that of the WT αM-II at 5 µM ATP. Interestingly, unlike WT myosin, MUT βM-II showed no effect on the crossbridge stiffness when ATP concentration was increased from 1 µM - 500 µM (p = 0.21 two tailed unpaired t-test) (Figure 7B). The observed high stiffness values ranging between 1.7 and 2 pN/nm for MUT myosins were comparable to the WT ventricular and atrial myosins observed at low ATP concentrations of 1 and 5 µM. For WT and MUT βM-II, we further confirmed that the state with a higher stiffness represents a ‘rigor-like’ state (Figure 7C and D). Here, instead of ATP, pyrophosphate (PPi) was used to attain primarily the strong-bound rigor-like AM state ^7, 29^. PPi was used in the reaction mixture to promote unbinding of the AM rigor complex, and high affinity of the PPi (10^7^M^-1^s^-1^) ^30^ enabled collection of individual binding/unbinding events. The binding events in the presence of PPi were analyzed and the stiffness was derived for single myosin molecules using variance-covariance method. As shown in Figure 7C, high stiffness of about 2 pN/nm was noted at 500 µM pyrophosphate, which was similar to the values obtained at 1 µM [ATP] for WT myosin (Figure 7D). Interestingly, for the MUT myosin, stiffness at 500 µM PPi was equivalent to the one at 500 µM ATP. The MUT and WT myosins shows comparable stiffness in the presence of PPi. Taken together, in contrast to WT myosin, the MUT myosin exhibited similar stiffness of ‘ADP-bound’ and ‘rigor’ state. The change of stiffness corresponding to the transition from ‘AM.ADP’ to ‘rigor’ state was not observed for the mutant myosins. It can be inferred that the MUT myosin may have overall high stiffness during active cross-bridge cycling at saturating ATP concentration under *in vivo* conditions.

### Reconstitution of human WT and MUT myosin

The 3-fold slower actomyosin detachment rate and shorter powerstroke size in single molecule measurements for the mutant myosin hinted at slower shortening speed (V = d / t_on_, where d-powerstroke size and t_on_- duration of AM attachment). Therefore, we wanted to examine the mutant myosins in an ensemble molecule actin filament gliding assay. As the remaining biopsy material was insufficient to extract required concentrations of myosins for the motility measurements, we employed alternative approach and reconstituted myosin with WT and MUT MLC1v.

Throughout this study, our objective was to employ native human cardiac myosins for the functionally closest comparison with the diseased motor composition in the patients. The LC1 reconstitution with pig, rabbit, and recombinant myosins have been reported earlier either with HMM or single headed myosin constructs ^17, 18, 31^. Here, for the very first time, we report reconstitution of the recombinant mutant MLC1v with human cardiac myosins obtained from donor tissue. The MLC1v was exchanged with a native full-length β-MII (Figure 8). Early reconstitution studies of MLC1v were proven to be challenging because of the significant loss of the myosin’s ATPase function. We further optimized the procedure by Wagner et al.^32^ to assemble MLC1v with the native human βM-II and obtained functional myosin (details in the SI). Note that to have a direct comparison between native myosins and reconstituted WT and MUT myosins, the tissue-derived myosin was subjected to the reconstitution procedure with WT and MUT MLC1v in parallel (schematic, Figure 8A). Up to 40% MLC1v exchange efficiency was achieved by mixing native myosin molecules with > 15-fold molar excess of MLC1v in a high-salt buffer at 15°C for 1h (Figure 8B and C).

**Figure 8.**
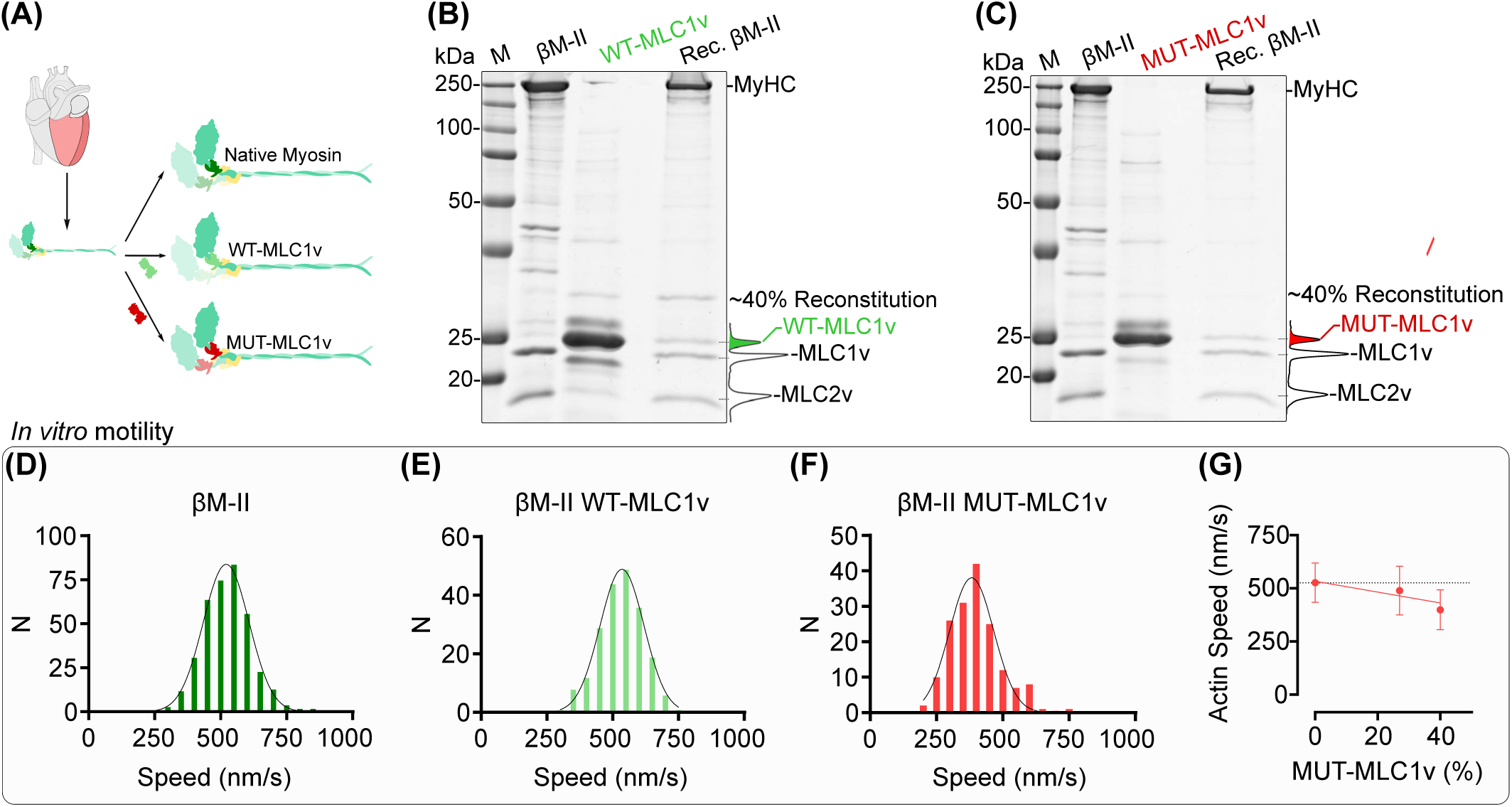
Reconstitution of human WT and MUT myosin. **A)** Schematic showing the extraction and reconstitution of myosin with the recombinant WT and MUT MLC1v. **B and C)** SDS-PAGE analysis of MLC1v reconstitution to human native ventricular myosin II (βM-II). βM-II reconstituted with recombinant WT-MLC1v (Rec. βM-II) with ∼40% efficiency, as determined by densitometry analysis employing the Gel Tool in ImageJ. A57D-MLC1v incorporation was similar (∼40%). MLC2v stayed intact during the reconstitution procedure. **D-F)** Actin filament gliding speed, fitted with Gaussian function. For native myosin, an average speed of 521.0 ± 86.9 nm/s (mean ± SD, n = 371) was comparable to that of WT-MLC1v reconstituted myosin, i.e., 535.0 ± 83.4 nm/s (p = 0.6153, nested t-test, n = 205). The filament speed distribution of the A57D-MLC1v reconstituted myosin had a mean speed of 382.7 ± 83.3 nm/s, significantly different from the native and WT-MLC1v reconstituted myosin (p = 0.005, nested t-test, n = 165). **G)** Mean actin filament speed plotted against the MLC1v reconstitution efficiency (%). Note that the data point for 0% is from native myosin. Solid line represents the linear regression fitting.

### MUT myosin shows reduced actin filament gliding speed

The native and reconstituted myosins were employed in the *in vitro* motility assay. The function of the myosin remained unperturbed following the exchange procedure as suggested by the comparable myosin-driven actin gliding speed for the native and WT-MLC1v reconstituted myosin, i.e., 521.0 ± 86.9 nm/s and 535.0 ± 83.4 nm/s, respectively (Figure 8D and E). Interestingly, A57D-MLC1v reconstituted myosin exhibited significantly slower actin gliding speed (Figure 8F). The reduction in the speed was proportional to the extent of reconstitution. The ∼27% reconstituted myosin displayed a gliding speed of 489.4 ± 114.8 nm, which decreased further to 398.7 ± 94.26 nm/s with ∼40% reconstitution (Figure 8G).

Altogether, the reconstituted WT βM-II mimicked the native full length βM-II function and the reconstituted MUT βM-II displayed slower actin filament gliding speed as expected from the changes in the velocity-determining parameters ‘t_on_’ and ‘d’ from our single-molecule measurements.

### Molecular dynamics simulations

To evaluate the change in structure-function relationship due to the A57D MLC1v mutation, we employed molecular dynamics simulations (MDS) and assessed the dynamic conformational changes underlying the experimentally observed mutation-induced changes in the AM dissociation rate, shortened powerstroke size and increased stiffness of the pliant region/lever arm.

### A57D modified the interaction of the MLC1v/ELC with the MyHC tail

To explore structural mechanisms though which A57D modifies crossbridge stiffness, we performed molecular dynamics (MD) simulations of a post-powerstroke actomyosin complex in AM.ADP* state (i.e., state 4 in Figure 9A.). Our simulations include a MyHC.ADP molecule in complex with an MLC1v, MLC2v, and f-actin pentamer (Figure 9B). In this chemomechanical state the upper and lower 50 kDa domains are engaged with actin, the nucleotide binding pocket is ‘open’, the N-terminal domain, converter domain, and tail are in the ‘down’ orientation. In the WT structure, A57 is located in the first α-helix of the MLC1v and is oriented towards a hydrophobic core of the MLC1v (Figure 9C). The A57 sidechain formed hydrophobic contacts with many MLC1v residues including F61, V79, and L83 as well as MyHC Y801 (Figure 9D and Table S1). The local structure around A57 was generally preserved in the WT simulation and maintained interactions with F61, V79, L83, and Y801. A57 remained sequestered from solvent: its average solvent accessible surface area (SASA) was 20 Å^2^ (∼9% of its total surface area). D57 became more solvent exposed: its average SASA was 50 Å^2^ (∼17% of its total surface area). D57 also maintained most native interactions, but its side chain formed a non-native hydrogen bond with MyHC residue Y801 (Figure 9D and Table S1). This structural change required repositioning of the N-terminal MLC1v α-helix.

**Figure 9.**
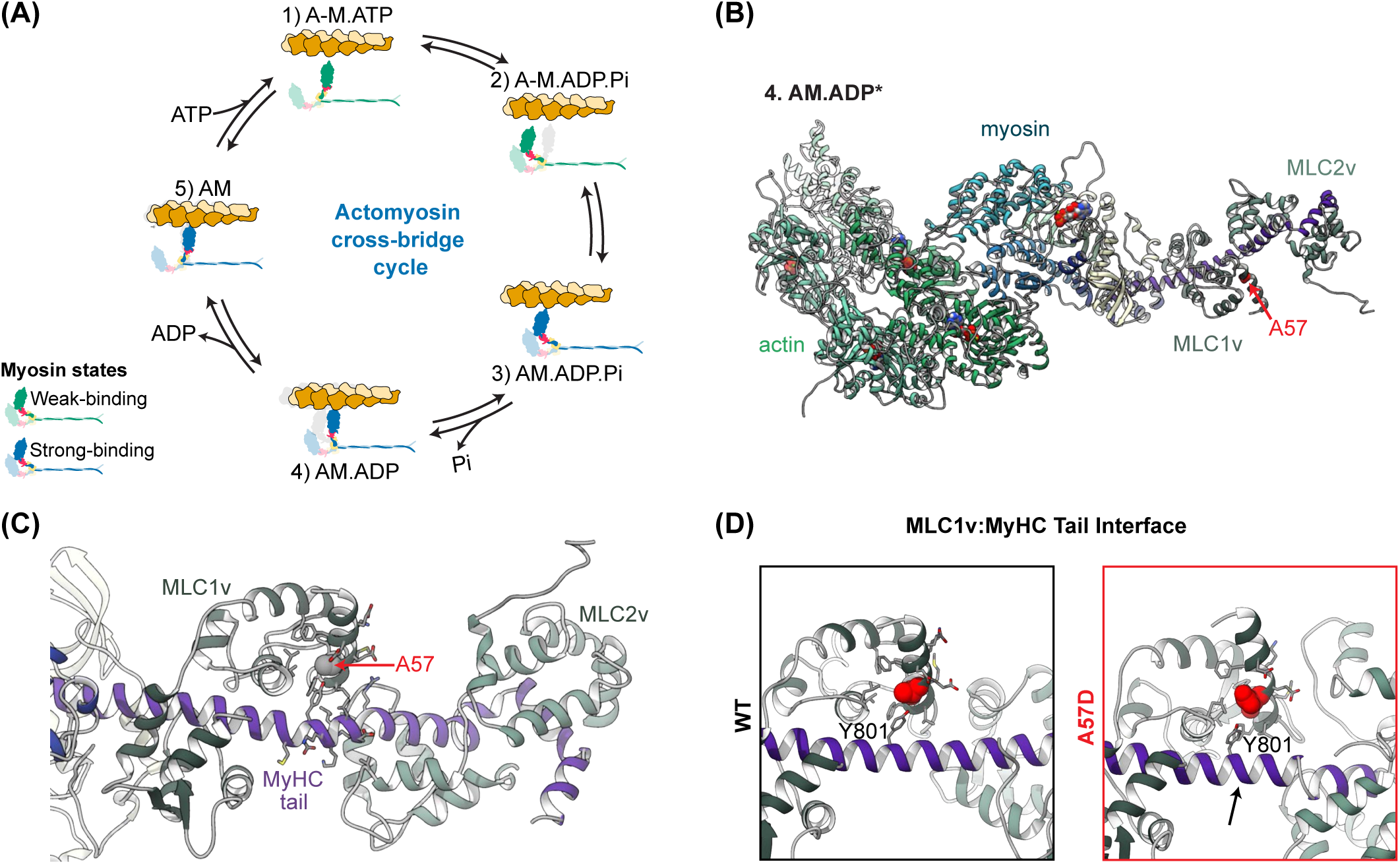
Molecular Models and Simulations. **(A)** This schematic summarizes the essential conformational states accessed during actomyosin crossbridge cycling. **(B)** The studied molecular model of the post-powerstroke state (4. AM.ADP*) includes an actin pentamer, myosin heavy chain, MLC1v, and MLC2v. Residue A57 is represented as red sphere. **(C)** The myosin pliant region with the essential and regulatory light chains bound to ‘IQ’ motifs within the myosin tail. The residue of interest in this study, A57 is situated in the hydrophobic core of the N-terminal lobe of the MLC1v. **(D)** Interaction between MyHC and MLC1v with A57 (WT) and D57 (MUT) MLC1v.

### A57D reduced the curvature and flexibility of myosin’s light chain binding region

The pliant region of the tail is involved in the myosin powerstroke, crossbridge flexibility, and formation of the interacting heads conformation of myosin. We quantified the dynamics of the tail by calculating the C_α_ root-mean-squared flexibility (RMSF), tail curvature, and hinge sites within the tail. We modeled the curvature of the tail by fitting Bezier curves to the backbone atoms of the pliant region. We found that A57D altered the curvature of the MyHC tail relative to WT (Figure 10A and B). In the A57D simulation, the portion of the tail region near the MLC1v binding site had ∼33% less curvature than was the WT simulation (Figure 10B). The repositioning of the MLC1v due to A57D also introduced a non-native kink into the pliant region around residue 810 which was not observed in the WT simulation (Figure 10C). There was also lower variance in the average curvature of residues 788-796 over time (Figure 10D). We also calculated the angle between two regions of the pliant region. The first angle was between residues 772-780 and 786-794 (where the MLC1v binds to the tail) (Figure 10E) and the second was between residues 777-789 and 802-817 (between the MLC1v and the MLC2v) (Figure 10F). For both angles, the WT simulation adopted more bent conformations and the A57D simulation adopted more linear conformations (Figure 10E and F). To analyze the average tail flexibility, we aligned our simulations on the C_α_ atoms of the pliant region (residues 767-823) to the average structure over the entire simulation and then calculated the C_α_ RMSF of tail residues, MLC1v residues, and MLC2v residues. The A57D simulation had reduced RMSF for every analyzed region (Figure 10G).

**Figure 10.**
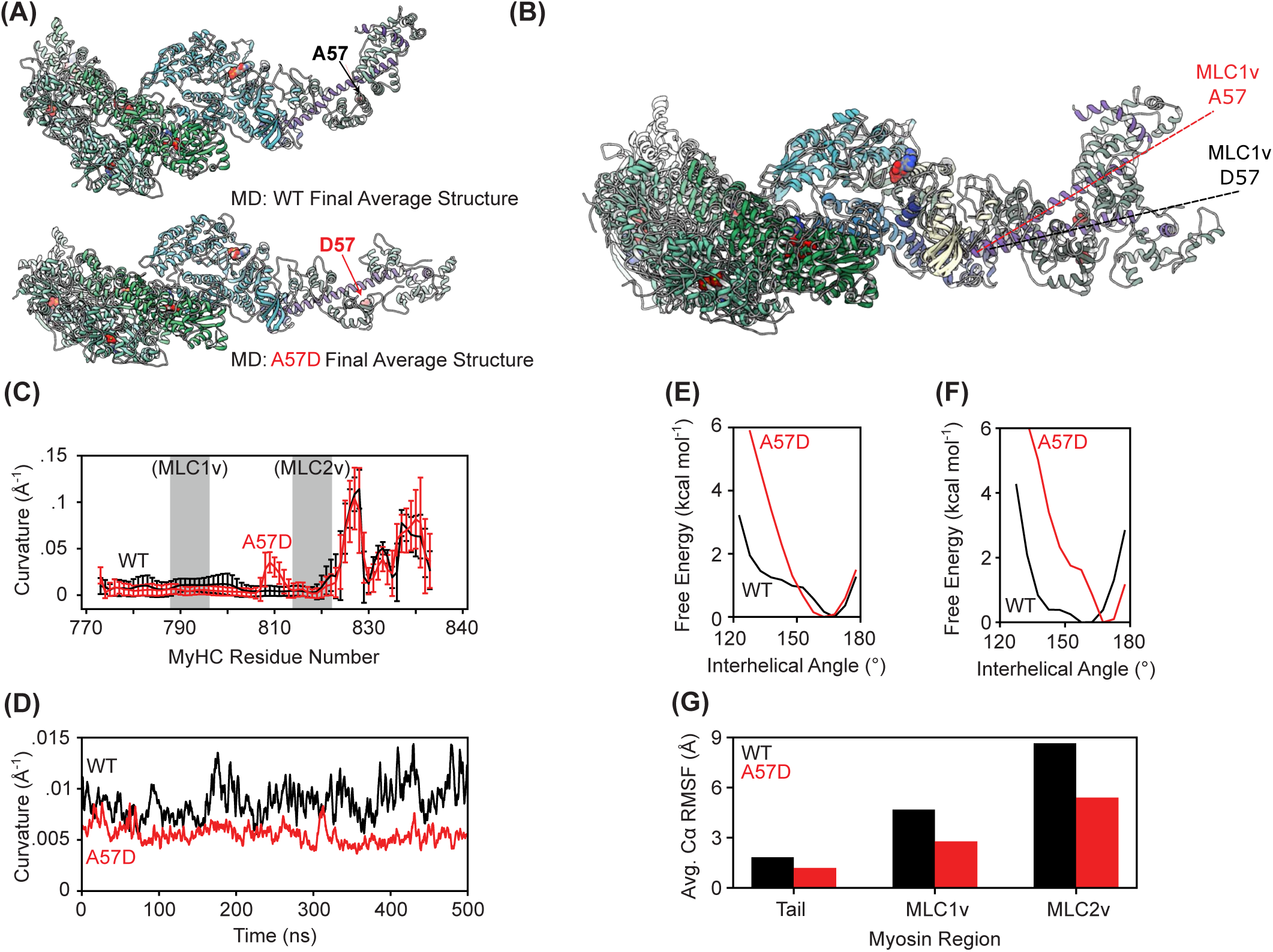
A57D modified the conformation of the MyHC pliant tail. **(A)** Final MD-derived conformations of the post-powerstroke actomyosin complex were generated by averaging over the final 50 ns of the A57 WT (top) and D57 MUT (bottom) simulations. **(B)** Superposition of the average final conformations of the WT and MUT simulations shows that the A57D mutation was associated with a change in the curvature of the myosin tail but not a change in the conformation of the motor domain. **(C)** Curvature of the MyHC tail as a function of pseudo-residue number for the WT (black) and A57D (red) simulations. The grey shaded regions denote the MLC1v and MLC2v binding sites. Error bars denote standard deviations after averaging across MD snapshots (temporal average). **(D)** The average curvature of the 33 N-terminal residues in the pliant region was plotted over time: the WT simulation not only had higher average curvature than did A57D but also had greater variance in the curvature. Interhelical angles measured within the MLC1v binding region of the tail **(E)** and in between the MLC1v and MLC2v binding sites **(F)** indicate that the WT simulation adopted more bent conformations than did A57D. **(G)** The simulations were aligned on the Cα atoms of the pliant tail and then the average RMSF values of the tail, MLC1v, and MLC2v were calculated. For all subregions analyzed the A57D simulation had reduced positional fluctuations relative to WT

### A57D allosterically altered the N-terminal domain of myosin and nucleotide binding pocket

Our molecular studies predict that the A57D mutation alters the interaction between the MLC1v and the MyHC tail in a way that alters tail curvature and flexibility (Figure 10 C and D). Changes in local structure due to A57D were also associated with altered MLC1v:MLC2v interactions and the interaction between the MLC1v and a helix in the N-terminal domain of myosin (Figure 11A). We also observed an altered conformation of ADP within the nucleotide binding pocket in which there were fewer interactions between ADP and the phosphate binding loop (residues 179-184) (Figure 11B). Prior studies on Omecamtiv Mercabil have noted an allosteric communication pathway that connects the N-terminal and converter domains to the nucleotide binding pocket ^33, 34^. Our simulations do not predict changes to the rest of the motor domain or major rearrangements of the actin-myosin interface (Figure 11C).

**Figure 11.**
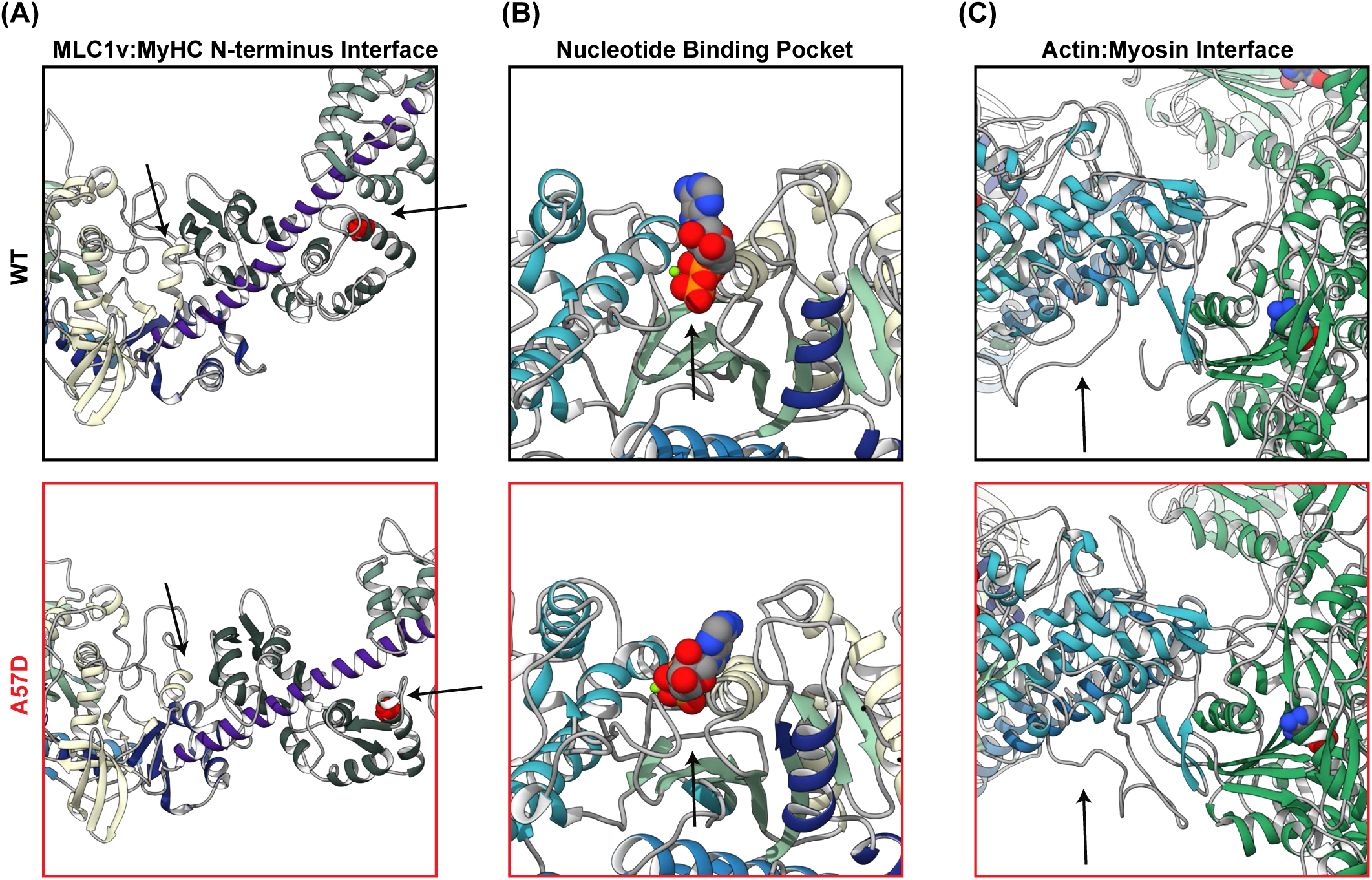
Summary of structural changes due to A57D. **(A)** Relative to WT, we observed that A57D led to local changes in the interactions between the MLC1v and the N-terminus of MyHC, **(B)** an alternate conformation of ADP within the nucleotide binding pocket, **(C)** and few changes at the actin-myosin interface. At the actin-myosin interface we did observe alternate conformations of the disordered loop 2 (indicated by an arrow).

Overall, molecular simulations predict that the A57D mutation causes a fundamental change in the residue-residue interaction network among the MLC1v, MLC2v, and MyHC. This altered interaction network resulted in a decrease in curvature and flexibility of the tail without affecting the length of the tail.

### Effect of A57D MLC1v on myofilament-level contractile properties

Computational modeling was used to predict how the molecular-level changes observed with A57D MLC1v MUT myosins affect the human cardiac muscle contraction. We employed FiberSim software ^35^, and modelled the sarcomere level changes in the contraction-relaxation parameters using experimentally derived powerstroke size (d) and detachment rate (k_cat_) as input parameters with either WT or MUT βM-II (Figure S2). The simulations of twitch response predict that muscle in A57D model exhibits a slower time-to-peak of 53.0 ± 3.6 ms compared to 38.7 ± 2.1 ms with WT model (Figure S2A). Additionally, a prolonged time from peak twitch force to 50% relaxation was predicted with a half relaxation time of 263 ± 1.7 ms for the A57D model compared to 180 ± 1 ms for the WT. Moreover, the simulations of force-pCa relationship predict an increased calcium sensitivity for A57D model with a pCa_50_ of 5.91 compared to 5.63 for the WT model (Figure S2B), whereby muscle with mutant myosins was predicted to generate a 34.2% higher force when comparing force at respective pCa_50_ calcium concentrations. Figure S2C and D shows single twitch kinetics and force-pCa relationship when only either powerstroke size or detachment rate was changed individually.

## Discussion

In this study, we investigated the impairment of primary cardiac myosin function caused by a point mutation in ventricular myosin light chain 1, responsible for pathological hypertrophic cardiac remodeling. We studied the biomechanical and biochemical mechanisms using a pure population of mutant motors from patient myocardium and wild-type motors from donor hearts, offering fundamental insights into the HCM pathology associated with the A57D MLC1v mutation.

### Primary motor dysfunction leads to pathological myocardial remodeling

As indicators of primary functional impairment of myosin, the major alterations in the chemomechanical cycle included: 1) changes in actomyosin cross-bridge kinetics, evident as a 3-fold reduction in actomyosin detachment rates, and 2) modification in mechanical properties, including a shorter powerstroke size and increased crossbridge stiffness (Table 1). These changes are predicted to decrease the contraction velocity. Consistent with this, analysis of reconstituted MUT βM-II containing the A57D MLC1v, demonstrated a reduced actin filament gliding speed compared to the WT βM-II. MD simulations revealed structural alterations in MLC1v and its modified interactions with MyHC and MLC2v, which may contribute to a stiffer lever arm, and conceivably, exert long-range allosteric effects on the biochemical and biomechanical properties of MUT motors.

**Table 1.** The effect of MLC1v A57D mutation on the kinetic and mechanical parameters derived from optical trapping measurements. *The power stroke size was estimated using ‘histogram shift’ method. 1^st^ and 2^nd^ powerstroke was estimated using ensemble-average method.

|  | Control<br>(WT MLC1v) | HCM<br>(MUT MLC1v) |
| --- | --- | --- |
| **AM detachment kinetics** |  |  |
| $k_{cat}$ (s<sup>-1</sup>) | 74 s<sup>-1</sup> | 26 s<sup>-1</sup> |
| $K_{0.5}$ | 34.9 μM | 13 μM |
| **Mechanical parameters** |  |  |
| **Powerstroke size (nm) \*** | 5.07 ± 0.19 nm | 3.4 ± 0.13 nm |
| **1<sup>st</sup> powerstroke** | 4.55 nm | 3.4 nm |
| **2<sup>nd</sup> powerstroke** | 0.6 nm | 0.23 nm |
| **AM Crossbridge Stiffness (pN/nm)** |  |  |
| **K<sub>1</sub> (ADP bound AM state)** | 0.62 ± 0.3 pN/nm | 1.70 ± 0.82 pN/nm |
| **K<sub>2</sub> (Rigor AM state)** | 1.92 ± 0.7 pN/nm | 1.99 ± 0.69 pN/nm |
| **Ensemble molecule assay (Reconstituted WT and MUT myosin)** |  |  |
| Actin filament gliding speed<br>at 40% exchange (nm/s) | 535 ± 83.4 | 398 ± 94.3 |

In addition to functional differences between WT and MUT motors, changes in myosin isoform composition were observed at the cellular level. Specifically, patient sample exhibited increased α-MyHC expression along with the corresponding light chain MLC1a. α-MyHC may assemble with ventricular subcomponents in combinations such as MLC1v-MLC2v or MLC1a-MLC2v, generating hybrid or chimeric motors, and thereby creating a heterogenous motor populations with diverse properties within a sarcomere. At the tissue-level, the diseased heart displayed cardiomyocyte disarray, interstitial fibrosis, and disrupted intercellular connections, with gap junction proteins (e.g., N-cadherin and connexins) losing their regular localization pattern. Coordinated cardiomyocyte contraction is a prerequisite for synchronous beating of the heart and mislocalization of gap junction proteins may impair cardiac conduction system. Furthermore, uncoordinated motor action and incoherent force production by a heterogeneous population of fast (α-MyHC) and slow MUT motor within the same sarcomere, as well as among different cardiomyocytes, may compromise overall mechanical output at organ level. These alterations likely activate function-compensatory pathways, including cardiomyocyte hypertrophy as indicated in the proposed model (Figure 12). Consequent cardiomyocyte disarray, expansion of myofibroblasts and activation of profibrotic signals promote the extracellular matrix (ECM) deposition, such as collagen, in the interstitial space, resulting in stiffening of the overall myocardium. Increased stiffness compromises the heart’s capacity to relax, as typically observed in HCM patients, contributing to diastolic dysfunction. In addition to fibrosis-induced stiffening, against which the cardiac tissue has to contract, the slower actomyosin dissociation kinetics observed in MUT motors may directly slow down ventricular relaxation, lowering preload and decreasing end-diastolic blood volume in the left ventricle.

**Figure 12.**
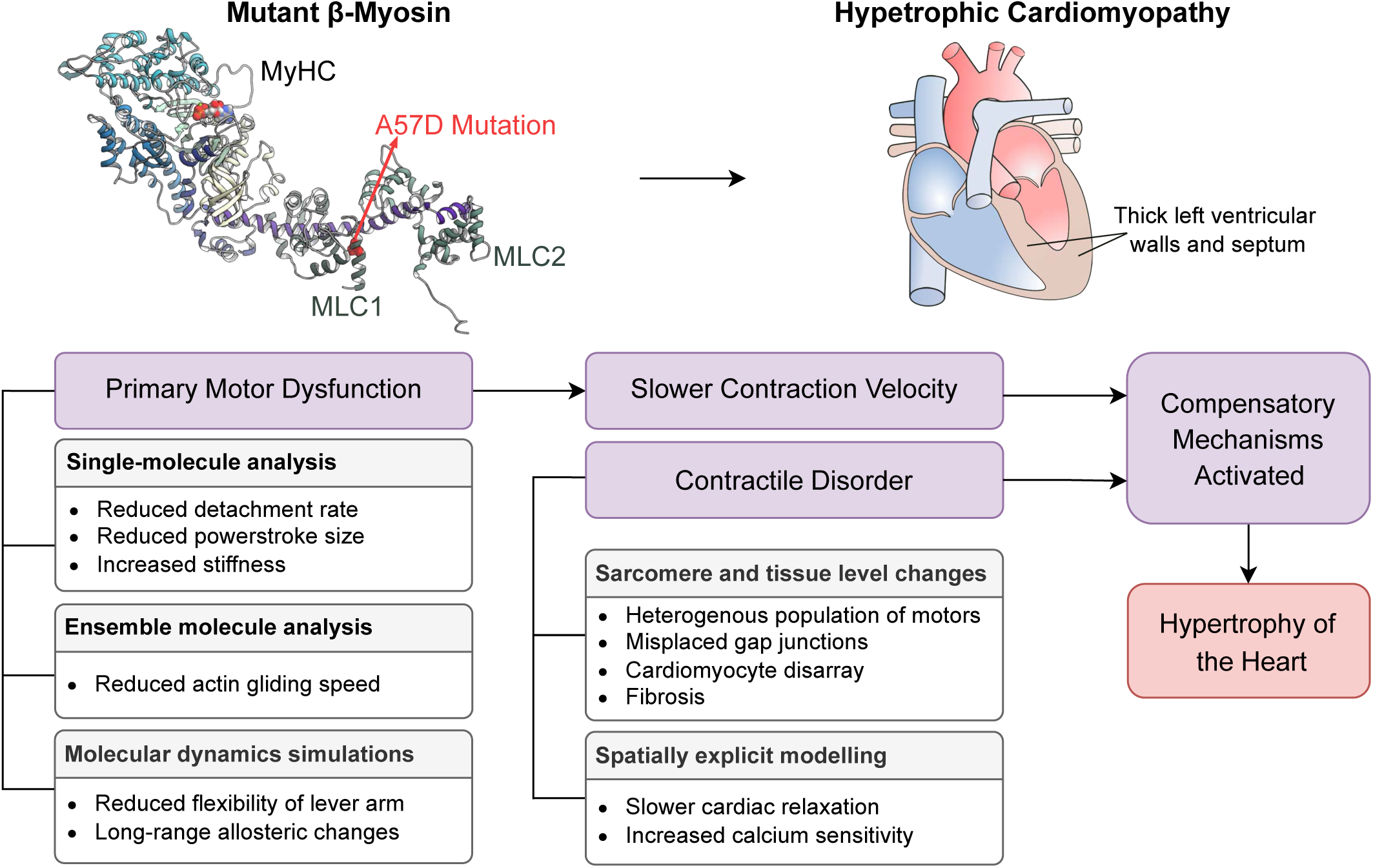
Proposed model. Summarizes experimentally observed alterations in the kinetic and mechanical properties of cardiac myosin, responsible for primary motor dysfunction at the single-molecule level and resulting in tissue-level changes. The possible path to pathological cardiac remodeling resulting from the MLC1v A57D mutation is outlined.

### Altered chemomechanical features as a consequence of MLC1v A57D mutation

Our single molecule analysis using trap assay indicate that the A57D MLC1v mutation increased the stiffness of actomyosin crossbridges in the post-powerstroke AM.ADP state. MD simulations predict that A57D modifies the structure of the lever arm in a way that reduces both the curvature and flexibility of the pliant region (Figure 10). It appears that this structural change in the MLC1v and thereby lever arm may be responsible for the changes observed in the chemomechanical states, i.e., the transition between the biochemical states from AM.ADP to AM state and reduced powerstroke size, through an allosteric effect that propagates through the molecule. We did not observe changes in the net length of the tail or in the actin-myosin interface due to the mutation, but we did observe an adjustment in the ADP binding to myosin (Figure 11). We earlier reported two actomyosin cross-bridge stiffness model ^7, 21^ whereby the AM.ADP state is more compliant (∼0.5 pN/nm) as compared to the AM rigor state (∼1.75 pN/nm). Interestingly, for the human WT βM-II, we found similar feature that the ADP bound AM state has lower stiffness than the rigor state. The measured values for stiffness of the two states were fairly comparable between the rabbit and human myosins, which also implies rather common feature among different myosin isoforms and species. Interestingly, this change in stiffness as the myosin progress through the crossbridge cycle was not observed for the MUT βM-II. The stiffness of the ADP bound state was same as the rigor state. There are two possibilities how the compliance of the AM bound states might influence the 2^nd^ powerstroke in MUT myosin, 1) lower ADP state stiffness compared to the rigor state maybe crucial for the generation of the second shorter powerstroke that is linked to the release of ADP from the active site, or 2) the stiffer pliant region may influence the full repriming of the lever arm thereby allowing partial swinging, and resulting in shorter powerstroke size. Overall, increased cross-bridge stiffness was observed for the AM.ADP force-generating state for MUT myosins. Furthermore, our simulations predicted mutation-induced changes in the curvature and flexibility of the pliant region, suggesting that these changes may increase the intrinsic stiffness of the pliant region and account for the increased stiffness observed experimentally.

Previous studies of HCM-causing mutations in the myosin converter domain, such as R719W and R723G, have demonstrated altered myosin head stiffness ^19, 36^, impacting the motile forces generated by myosin. These findings led to the conclusion that the converter region is the primary elastic element within the myosin motor domain. Our current findings with the MLC1v mutation, which also results in increased stiffness, suggest that the lever arm domain and its associated light chains serve as an additional, instrumental elastic element directly modulating actomyosin cross-bridge stiffness.

Recently, the relevance of fraction of myosin heads in the sequestered or inhibited state *vs.* the weak binding or disordered states has gained significant attention for cardiac function and as a means for therapeutic intervention in specific HCM conditions ^37, 38^. A number of HCM carrying mutations in MyHC have been linked to increased fraction of myosin heads in the disordered state and thereby generating higher forces in comparison to wild type myosins ^38–40^. As conformation and dynamics of the tail affect stability of sequestered heads, altered myosin recruitment may also be an effect of A57D. Although not directly assessed for A57D MLC1v mutation here, conceivably, the additional interactions between the MyHC (Y801) and D57 may impact the stability of the sequestered state of myosin by increasing the energetic barrier to the formation of the interacting heads motif by reducing flexibility at hinge sites necessary for the heads to interact with one another. Our simulations do not account for the structure and dynamics of the tail within a sarcomere or crowding effects that would be present in two headed myosin constructs, since our model construct does not include density for S2 and we have simulated only a single motor domain bound to an actin pentamer. Besides, our simulated crossbridges are not under tension. Modeling efforts may be improved in the future by considering the effects of increased crossbridge stiffness in a more complex molecular environment. Nevertheless, the increased cross-bridge stiffness is expected to affect the force generation.

Mechanical work done *(W)* per ATP hydrolyzed can be estimated using powerstroke size and stiffness values from our single molecule measurements. Using a function for the potential energy of a spring, i.e., 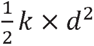 (*d* - overall stroke size/displacement, *k –stiffness*/spring constant), we calculated the mechanical work done per ATP molecule per AM crossbridge cycle. Using overall stroke size of 5.07 nm and 3.4 nm and stiffness values of 1.93 and 2 pN/nm for WT and MUT βM-II, respectively, the work done were calculated to be 24.8 and 11.56 pN.nm, respectively. Note that here we used the rigor stiffness of the AM crossbridges, i.e., the state when the powerstroke is completed. Thus, the MUT motors perform nearly two times lower mechanical work per ATP hydrolysis. From the ATP concentration dependence of the actomyosin detachment rates, we further estimated the 2^nd^ order rate constant for ATP binding to the WT and MUT βM-II from linear fitting of the initial measured rates. These values were comparable with 1.75 ± 0.08 µM^-1^s^-1^ and 1.31 ±0.1 µM^-1^s^-1^ for WT and MUT motors, respectively, implying that the ADP release is the main rate limiting step that reduces the actin filament gliding speed in MUT myosin.

### Pure native mutant motors to assess the direct effects of mutation

In this study, we opted for homogeneous population of human cardiac full-length native myosins to investigate their function, which is directly relevant for human physiology and pathophysiology. Previous reports using patient-derived native myosins-either from M. soleus or cardiac tissue of individuals carrying HCM-associated MyHC mutations (R403Q and L908V) - demonstrated significantly reduced actin filament sliding velocities ^41, 42^. Affected individuals were heterozygous for these mutations, and thereby expressed variable amounts of WT and MUT motors. Consequently, the observed effects on sliding speed reflected the mixture of WT and MUT myosins, complicating the determination of mutation-specific contributions from the ensemble measurements. Moreover, studies involving myosin or myofibrils derived from alternative tissue sources need to be carefully evaluated for tissue-specific differences. We previously demonstrated that the βM-II isoform from cardiac tissue exhibits distinct mechanical properties compared to M. soleus-derived myosin motors, despite both expressing the same β-myosin heavy chain isoform (β-MyHCs) ^7^. This suggests that the kinetic and mechanical effects studied using soleus muscle cannot be directly extrapolated to cardiac function. Studies on transgenic animal models further revealed variations arising from species-specific differences, for example the R403Q MyHC mutation showed increased actin-activated ATPase activity and actin filament sliding speed in cardiac samples of transgenic HCM mice, while transgenic rabbit myosin with the same mutation showed reduced actin-activated ATPase activity and actin sliding speed that is more consistent with patient tissue-derived myosin ^43, 44^. The discrepancies likely arose from difference in the MyHC backbone, i.e., α-MyHC in mouse vs β-MyHC in rabbit. The composition of endogenous light chains partners further adds complexity, as the light chains modulate heavy chain function and can influence the mechanical and kinetic properties, as reported earlier for different myosins ^7, 21, 45, 46^.

In recent years, expression of human cardiac MyHC in murine myotubes (C2C12) has enabled the investigation of MyHC isoform-specific differences, assessment of numerous disease-causing mutations for their functional impairments, elucidation of structure-function relationships, and facilitation of screening and testing of cardiomyopathy drugs ^33, 38, 47–49^. Some of these studies have utilized truncated recombinant beta-cardiac myosins - either single headed myosin subfragment-1 (S1, comprising approximately 808 - 845 aa of MyHC) or double-headed heavy meromyosin (HMM) – in which the associated light chains (LCs) were either of murine origin, in some cases only human ELC, and in fewer cases both of human origin ^38^. Truncated myosin constructs, which lack the long tail domains, remain a preferred choice for certain experimental settings due to the challenges associated with the stability and limited experimental adaptability of full-length myosins. Nevertheless, differences in the composition of the myosin complex may directly influence its function and, consequently, the findings. Our studies emphasize the need to consider species- and tissue-specific differences, when examining human cardiac disorders.

### Advantages of reconstitution approach

The limited availability of human heart samples from affected patients presents significant challenges for studying HCM. We addressed this limitation by establishing and optimizing a reconstitution system that utilizes human native myosin from small donor samples, in combination with recombinant WT MLC1v and MUT A57D-MLC1v. While MLC1v exchange has been conducted in pig myosins ^18^, it has not previously been reported in human myosins. Our approach enables direct comparison of endogenous myosins with reconstituted human WT and MUT myosins, thus providing a clinically relevant assessment, and eliminating concerns about isoform-, and species-specific differences in key motor function parameters. Additionally, the ability to control exchange efficiency allows generation and investigation of heterozygous mixture of WT and MUT myosins, closely mimicking the myosin composition found in many patients and thought to contribute to sarcomeric and myofibrillar disarray.

The mechanistic understanding of many HCM-causing mutations in the *MYL3* gene remains rather limited. Some MLC1v mutations have been investigated using transgenic animal models ^9, 50^. A mutation at the same residue as our studied variant, i.e., A57G in MLC1v, has been reported to cause a particularly severe form of HCM. Examination of cardiac muscles from transgenic mice carrying the MLC1v A57G mutation revealed a significant increase in mutation-induced Ca²⁺ sensitivity of force generation, about 1.3-fold reduction in maximal force, and increased myocardial stiffness ^17^. With our successful reconstitution approach, extending these findings in mouse cardiac tissue to human-derived one will be important for drawing parallels to human cardiomyopathy in the future. Collectively, validation of this human myosin reconstitution opens new avenues for mechanistic studies of unexplored mutations and provides a platform to evaluate the efficacy of potential drugs directly applicable to human myocardium.

Our findings on the effects of A57D MLC1v mutations at the molecular level reveal the underlying mechanisms that trigger the development of the HCM phenotype. Diastolic dysfunction is a characteristic signature of HCM. Mechanistically, we demonstrated slowed actomyosin detachment kinetics for the MUT motor, as a contributing factor to impaired cardiac relaxation, as predicted by simulations of sarcomere-level changes in cardiomyocytes (Figure S2). The pronounced impact of MLC1v mutation on βM-II function exemplifies myosin’s long-range allosteric communication. Importantly, the ability to reconstitute human full-length native myosin with chosen LC mutations that mimic the tissue-derived mutant proteins provide a valuable approach to study these mutations without reliance on patient samples. This study underscores the importance of examining individual mutated motor components in their correct molecular context to gain detailed insights into the underlying pathomechanisms – an essential step toward designing targeted therapeutic strategies.

## Methods

### Human myocardium samples

Human myocardial sample was obtained from a patient with hypertrophic cardiomyopathy undergoing surgical septal myectomy for relief of left ventricular outflow tract obstruction (LVOTO). Non-diseased, non-transplanted heart muscle tissue or donor samples were obtained from Sydney heart bank, Australia. Local ethical approval was obtained from Hannover Medical School ethics committees for use of tissue samples (No. 1749-2013). According to the Declaration of Helsinki (20), written informed consent was given by subject.

### Patient and donor cardiac tissue samples

A 52-year-old male patient with hypertrophic obstructive cardiomyopathy (HOCM) underwent a myectomy procedure. The patient harbored a homozygous mutation c.170C>A in the *MYL3* gene that encodes ventricular myosin light chain 1 (MLC1v). The excised tissue sample was employed to study the impact of the homozygous mutation A57D in MLC1v on β-cardiac myosin (βM-II) function. Age and gender-matched left ventricular tissue sample from a 56-year-old male was selected as a control wild type (WT) for comparison with the above mutant. The tissue samples were immediately flash-frozen in liquid nitrogen after myectomy for a patient or harvested post-death from a non-diseased human heart. The storage procedure was adjusted to minimize the influence of tissue handling on the protein quality and to preserve the native function of the biomolecules. To optimize the usage of the small-size human tissue sample for various experiments, we adopted the workflow schematized in Figure S1.

### Immunohistochemistry

To obtain cryosections the tissue was embedded in Tissue-Tek O.C.T. (Sakure Finetek Europe, Leiden, NL) on typical specimen stages at -20°C in a cryotome (Leica CM1860 UV, Wetzlar, Germany). Single cryosections of 5 µm thickness were attached to HistoBond adhesive microscope slides (Marienfeld GmbH & Co.KG, Lauda-Königshofen, Germany) and immediately fixed in 4% PFA (paraformaldehyde, Merck KGaA, Darmstadt, Germany) for one hour at room temperature. After washing with PBS (Dulbecco, Biochrom GmbH, Berlin, Germany) the tissue slices were permeabilized in 0.5% Triton X-100 (Roche, Basel, Switzerland) and followed by immunofluorescent labelling with primary antibodies for one hour. After washing out the primary antibody using PBS, the cryosections were incubated with secondary antibodies for another hour. DAPI (D9542, Sigma-Aldrich, Deisenhofen, Germany) was used to stain nuclei. In the end Fluoroshield (Sigma-Aldrich, Deisenhofen, Germany) was applied to cover the tissue between microscope slide and a small glass coverslip. The slides were visualized using the Epifluorescence microscope.

### List of primary and secondary antibodies

| Primary Antibodies |  |  |  |  |
| --- | --- | --- | --- | --- |
| Anti-Collagen1 | 1:200 | from mouse | Thermo MA1-26771 | Alexa 555 (d anti m) |
| Anti-αActinin | 1:200 | from rabbit | Abcam ab137346 | Alexa 488 (d anti rb) |
| Anti-nCadherin | 1:200 | from goat | ABIN 1440032 | Alexa 680 (d anti g) |
| Anti-β-MyHC | 1:100 | from mouse | Sigma M8421 | Alexa 555 (d anti m) |
| Anti-αMyHC | 1:50 | from rabbit | rabbit-α-huMYH6, polyclonal, BioGenes, Berlin | Alexa 488 (d anti rb) |
| Anti-Connexin43 | 1:200 | from mouse | Merck MAB3067 | Alexa 555 (d anti m) |
| Secondary antibodies |  |  |  |  |
| Alexa 555 (d anti m) | 1:400 | Life A31570 | Anti-Collagen1, Anti-β-MyHC, Anti-Connexin43 | |
| Alexa 488 (d anti rb) | 1:400 | Life A21206 | Anti-αActinin, Anti-αMyHC | |
| Alexa 680 (d anti g) | 1:400 | Life A32860 | Anti-nCadherin |  |

#### Native myosin extraction from human heart muscle

Human cardiac tissue was crushed using a liquid-nitrogen-cooled mortar and pestle. The resulting powder was aliquoted and stored in -150°C. To extract the myosin, high salt extraction buffer (0.5 M NaCl, 10 mM HEPES, pH, 7.0, 5 mM MgCl_2_, 2.5 mM MgATP) was used. Typically, to one ∼ 10 mg aliquot of tissue powder, an ice-cold extraction buffer supplemented with 2 mM DTT and 0.5 mM AEBSF was added at 1:5 (w/v) ratio ^51, 52^. The myosin was extracted at 4°C for 20 min on ice and then centrifuged for 45 min at 65000 rpm. The supernatant was collected and diluted 10 fold with ice-cold water containing 2 mM DTT and allowed to precipitate on ice for 40 minutes. It was then spun for 30 min at 27,000 rpm at 4°C to collect the precipitated myosin. The pellet was dissolved in myosin extraction buffer with 2 mM DTT and 0.5 mM AEBSF. The concentration of myosin was determined using the Bradford assay and the purity was assessed using SDS –PAGE as shown in Figure 4. Isolated myosin was aliquoted, flash frozen in liquid nitrogen and stored at -80°C in 50% glycerol.

#### Preparation of actin filaments

Actin was purified from chicken pectoralis muscle as described ^53^. To obtain sufficiently longer biotinylated actin filaments (≥ 20 µm) for optical trapping experiments, chicken G actin and biotinylated G actin was mixed in equimolar ratios to a final concentration of 0.1 µg/µl each in p-buffer (5 mM Na-phosphate, 50 mM K-acetate, and 2 mM Mg-acetate), containing 1 mM DTT, 1 mM ATP, and 0.5 mM AEBSF protease inhibitor (Cat No. 30827-99-7, PanReac Applichem ITW). The mixture was incubated overnight at 4°C, and followed by addition of equimolar concentration of fluorescent (TMR, tertramethylrhodamine) phalloidin (cat no. P1951, Sigma-Aldrich) and biotin phalloidin (0.23 nM, Invitrogen/Thermofischer Scientific, B7474) to label the actin filaments.

#### 3-bead assay using optical trap

The optical trapping set up was described in detail previously ^54, 55^. For the assay, flow cells with approximately 15 µl chamber volumes were assembled using coverslips with nitrocellulose-coated beads. Glass microspheres (1.5 µm) suspended in 0.05 % nitrocellulose in amyl acetate were applied to 18x18 mm coverslips. All the dilutions of biotin-actin filaments were made in reaction buffer (KS buffer) containing 25 mM KCl, 25 mM HEPES (pH 7.4), 4 mM MgCl_2_, and 1 mM DTT. The full-length native myosin was diluted in high salt extraction buffer without MgATP. For the experiment, the chamber was prepared as follows, 1) flow cells were first incubated with 1 µg/ml native myosin for 1 min, 2) washed with high salt extraction buffer without ATP and thereafter with KS buffer, 3) followed by wash with 1 mg/ml BSA and incubated further for 2 min to block the surface, 4) finally, reaction mixture containing 0.8 µm neutravidin coated polystyrene beads (Polyscience, USA) and 1-2 nM biotinylated actin was flowed in with 5 µMATP (or varied concentrations of ATP), ATP regenerating system (10 mM creatine phosphate, and 0.01 unit creating kinase) and deoxygenating system (0.2 mg/ml catalase, 0.8 mg/ml glucose oxidase, 2 mg/ml glucose, and 20 mM DTT). The assembled flow chamber was sealed with silicon and placed on an inverted microscope for imaging and trapping assay.

An actin filament was suspended in between the two laser trapped beads (Figure 4A), pre-stretched, and brought in contact with the platform bead immobilized on the chamber surface. Low-compliance links between the trapped beads and the filament were adjusted to about 0.2 pN/nm or higher ^56^. The bead positions were precisely detected with two 4-quadrant photodetectors (QD), recorded and analyzed. The actomyosin interaction events were monitored as a reduction in free Brownian noise of the two trapped beads. Data traces were collected at a sampling rate of 10,000 Hz and low-pass filtered at 5000 Hz. All the experiments were carried out at room temperature of approximately 22°C.

#### AM crossbridge stiffness measurement and analysis

The AM interactions were recorded by applying positive position-feedback on the laser-trapped beads as described in Steffen et al ^7, 21, 54, 57^. We used analog positive feedback in AC mode to increase the noise amplitude of the free dumbbell. Consequently, the variance ratio between free dumbbell noise and binding events improved for both traps in the direction of the actin filament axis (x-axis). This practice allowed event detections and stiffness measurements by increasing the amplitude of Brownian noise by ∼30%. Two different methods were employed to measure the stiffness (k) of the myosin motors, namely – ‘variance-covariance’ method and ‘ramp’ method as described in detail previously ^56, 58^. Using the variance-Hidden-Markov-method, acto-myosin interactions events detected as reduction in noise were analyzed. The position variance-covariance of two trapped beads in both the bound and unbound states of actomyosin interaction offers an alternative means to measure the myosin stiffness. The data records with combined trap stiffness in the range 0.06–0.09 pN/nm were used. Low-compliance links between the trapped beads and the filament were adjusted such that the ratio of the position variance during free and bound periods was in the range of 3-10 as described in Smith et al.^58^. Due to limitations in resolution of optical trap assay and subsequent difficulty in reliable molecular event detection, the WT ßM-II was assessed at a maximal ATP concentration of 50 μM. In contrast, the MUT ßM-II permitted the use of a higher concentration, up to 500 μM ATP. To establish a comparative reference for the ‘rigor-like’ state stiffness, independent measurements were performed at 500 μM pyrophosphate (PPi) in KS buffer.

Briefly, for the ‘ramp’ method, a big triangular wave was applied on both the beads. A large amplitude triangular wave of 240 nm and 1 Hz was chosen to study acto-myosin binding events at low ATP concentration (5 μM) due to their long life times, while 120 nm amplitude and 2 Hz wave was applied at ATP concentrations of 50 µM ATP for WT ßM-II and 100 μM ATP for MUT ßM-II. A constant ramp velocity of 480 nm/s was used for high and low ATP concentrations. The AM interaction events were registered on both upwards or downwards sides of the ramp. For the AM binding events, the velocity ratios between unbound and bound states for left and right beads were calculated. The AM cross-bridge stiffness was deduced using the velocity ratios, trap stiffness and combined link stiffness as described in details previously ^21, 56^.

#### In vitro motility assay

The details are provided in the supplemental information

## Data availability

All the data supporting the findings of this study are available within the article, the supplementary information files, and the source data files.

## General

We are grateful to Petra Uta for the technical assistance in protein preparations, and Ante Radocaj for critical comments on the manuscript.

## Author contributions

MA and AN conceived the project. MA designed the experiments with inputs from TW, ES and AN. TW performed optical trapping measurements and analyzed the data. ES performed reconstitution and motility experiments and analyzed the data. MC and MR were involved in molecular dynamic simulations. TH performed the immunohistochemistry. KC and ES performed the FiberSim modelling. CDR provided the donor heart tissue samples. TT provided patient tissue. TK was involved in the discussions. MA supervised the project and wrote the manuscript with assistance from TW, ES, MC, and AN. All the authors contributed to the editing of the manuscript.

