## Supplementary Information for "Single-molecule analysis sheds light on cardiac myosin dysfunction due to hypertrophic cardiomyopathy mutation A57D in ventricular myosin light chain-1 (MLC1v)"

**Methods –**

**Mutagenesis and expression of human MLC1v**

The expression vector containing the human ventricular myosin essential light chain insert (i.e., MYL3) with an N-His tag was purchased from Genecopoeia (cat# EX-D0092-B11-GV0-GC). Site-directed mutagenesis was employed to introduce the Ala57Asp (A57D) mutation. Template and mutant expression vector sequences were confirmed using Sanger sequencing to compare with the expected reference sequence (NM_000258.3), and were used to transform Rosetta (DE3) pLysS cells (Sigma-Aldrich, 70956) for the expression of WT- and A57D-MLC1v proteins. The transformed cells were grown overnight at 37 °C spread on 50 µg/ml Ampicillin-containing agar plates. Individual colonies were grown overnight in 20 ml Terrific Broth media (TB) at 25 °C and 150 rpm shaking speed. The following day, cells were transferred to Erlenmeyer flasks containing 200 ml Terrific Broth. When the culture reached the optimal density of $OD_{600} \approx 0.6$, protein expression was induced using 1 mM IPTG (Sigma-Aldrich, I6758) and cells were incubated overnight at 16 °C and 120 rpm orbital shaker speed. Cells were harvested the next day by centrifugation at 8000 rpm for 10 min at 4 °C (JLA 10.500 rotor). The resulting cell pellet was washed by resuspending in 20 ml cold DPBS and centrifuged for 10 min at 4000 rpm (Biofuge Hereaus 7588 rotor). The washed pellets were stored at -80 °C until purification.

**Purification of MLC1v**

Each cell pellet was thawed and resuspended in 10 ml lysis buffer (40 mM Imidazole, 500 mM NaCl, 50 mM NaH2PO4, 1 mM EDTA, 10 % Glycerol, pH 8.0 at 4 °C) supplemented with 1 mM DTT, 1 mg/ml lysozyme, 1 % TritonX-100, 1 mM AEBSF, 1x cOmplete protease inhibitor cocktail (PIC), and 5 mM MgATP for 10 min at 37 °C, followed by the addition of 10 µg/ml of DNAse I and 20 mM $MgCl_2$ and incubation for additional 10 min at 37 °C. Next, the lysate was sonicated on ice for two cycles of 1 min 40 s (40% amplitude, 20 s pulse, 40 s pause), incubated for 30 min at 4 °C, and centrifuged at 10,000 g for 30 min at 4 °C (Biofuge Hereaus 7588 rotor).

The clarified lysate was subjected to nickel affinity purification to isolate the HIS-tagged MLC1v protein. A 20 ml gravity flow column was loaded with 1 ml of 50% suspension PureCube 100 Ni-INDIGO Agarose nickel resin (Cube Biotech, 75103). Resin was equilibrated using lysis buffer, followed by incubation with 10 ml cleared lysate on a rotator for 1 h at 4 °C. The resin was washed with four cycles of 10 ml wash buffer (same as lysis buffer but with 50 mM Imidazole and without lysozyme, TritonX-100, and DNAse I). The bound HIS-tagged MLC1v protein was eluted four times using high imidazole elution buffer (500 mM Imidazole buffer containing 300 mM NaCl, 50 mM NaH2PO4, 1 mM EDTA, supplemented with fresh 1 mM DTT, and 1 mM AEBSF) and ~5-15 min incubation. The buffer of combined eluate fractions was exchanged into wash buffer by employing 10 kDa cut-off 15 ml Amicon spin columns. The purified protein was stored at -80 °C.

**Myosin purification and reconstitution with expressed MLC1v**

The native myosin was purified using the protocol published previously[1], with small modifications. Briefly, tissue samples were ground into small pieces in liquid nitrogen using a pestle and mortar, followed by the collection of the pieces into an ultracentrifuge tube containing a 4:1 (v/w) ratio of pre-chilled myosin extraction buffer (500 mM NaCl, 10 mM HEPES pH 7.0, 5 mM $MgCl_2$, 2.5 mM MgATP, supplemented with fresh 2 mM DTT, 1 mM AEBSF, 1x PIC) and 20 min incubation on ice with intermittent mixes. The mixture was centrifuged for 60 min (4 °C, 150 000g, TLA120.2) to obtain the crude myosin extract, which was divided into two ultracentrifuge tubes for further purification; one tube containing $1/5^{th}$ of the total myosin volume ("1x myosin pellet"), and the other tube containing $4/5^{th}$ of the volume, the "4x myosin pellet". Myosin in both tubes was diluted 10x (v/v) with ultrapure water containing 2 mM DTT, and incubated for 40 min on ice to reduce the ionic strength and promote myosin filament formation. Myosin filaments were collected by a 40 min centrifugation (70 000 g, 4 °C, TLA 110). The "1x myosin pellet" was used for Bradford protein quantification and myosin molarity calculation, with the remaining serving as native myosin control. The "4x myosin pellet" was used for the MLC1v reconstitution procedure.

MLC1v reconstitution protocol was based on Wagner and Weeds[2], with some modifications. In principle, a reconstitution mixture with > 15-fold excess of MLC1v to myosin was aimed. For this, the purified MLC1v aliquots from -80 °C were thawed on ice, and 0.5 ml 10-kDa Amicon spin columns were used to exchange the buffer into MLC1v reconstitution buffer (100 mM Imidazole, 4.7 M $NH_4Cl$, 2 mM EDTA, supplemented with 2 mM DTT, 1 mM AEBSF, 4 mM $MgCl_2$, and 2 mM MgATP). The MLC1v concentration after buffer exchange was determined by Bradford assay, and a correction factor was applied to account for the purity of the MLC1v band (~50-60%) in the Coomassie-stained SDS-PAGE documented after the MLC1v purification procedure, followed by molarity calculation (1 mg/ml sample of MLC1v corresponding to 45.59 µM). The calculated amount of MLC1v was then used to dissolve the "4x myosin pellet", followed by an incubation for 1 h at 15 °C using a thermal shaker set at 900 rpm. The reconstitution mixture was then loaded on a 0.5 ml 100 kDa Amicon spin column to remove unbound MLC1v, followed by a 30x dilution using ultrapure water containing 2 mM DTT. This sample was incubated for 40 min on ice to ensure myosin filament formation, and subsequent 40 min centrifugation was used to precipitate the myosin filaments (70 000 g, 4 °C, TLA 110). The final myosin pellet was dissolved using myosin extraction buffer, and was either used fresh in functional studies or flash-frozen by pipetting 10 µl directly into liquid nitrogen, forming small-sized beads to be stored at -80 °C.

Reconstitution efficiency was assessed by densitometric analysis of SDS-PAGE gels using ImageJ.

### *In vitro* motility assay

Actin gliding assay was used for the functional assessment of the WT- and A57D-MLC1v reconstituted myosin, performed as explained before[1] with minor modifications. Briefly, ~5 µl flow chambers were prepared and covered with BSA, followed by myosin infusion and BSA blocking. Next, non-functional myosin motors were blocked with unlabeled actin, followed by the infusion of rhodamine-phalloidin labeled actin filaments. The movement of labelled actin filaments was started using the assay buffer (25 mM imidazole pH 7.2, 25 mM NaCl, 4 mM $MgCl_2$, 1 mM EGTA, 2 mM DTT, 0.5% methylcellulose), containing the anti-bleeching system (10 mM DTT, 18 µg/ml catalase, 0.08 mg/ml glucose oxidase, 10 mg/ml D-Glucose). Movies were aqcuired using a custom-made TIRF microscope at 22 °C for 300 frames at 5 fps with 170 nm pixel resolution. Smoothly moving filaments were manually tracked using the MtrackJ plugin[3] in ImageJ.

### Molecular Modeling and System Preparation

We constructed our model of the post-powerstroke, force-bearing actomyosin complex (Figure 9 A-B, 4.AM.ADP*) using the cryoEM structures of ADP-state myosin + ELC and the actin-tropomyosin-myosin complex presented by Doran *et al.* (PDB IDs 8EFE[4,5] and 8EFH[5,6]). The latter structure was solved via helical reconstruction and was missing atomic density for the pliant tail and essential light chain. Coordinates corresponding to myosin heavy chain (MyHC) from residues 766-796 and ELC residues 46-188 were transferred from 8EFE to 8EFH. Next, we removed the tropomyosin strand. The thin filament was modeled as an actin pentamer in the absence of tropomyosin: three actin subunits formed the protofilament that was bound to myosin and two subunits formed the other protofilament. Finally, missing coordinates for the light chain binding region of the tail MyHC (gene: myh7, Uniprot Entry: P12883) residues 766-850, ELC (gene: *myl3*, Uniprot Entry: P08590) residues 46-195, and RLC residues 1-166 (gene: myl2, Uniprot Entry: P10916) were built using homology modeling (*Modeller*[7]). The structure of the scallop adductor muscle myosin II light chain binding region in complex with its associated ELC and RLC (PDB: 3TS5 [8,9]) was used as a template for modeling the tail and RLC. This modeling strategy makes several assumptions, and our overall model is limited by the fact that simulations of the human system were initiated from a composite structure. The limitations are as follows. First, the disordered N-terminal extension of cardiac ELC (residues 1-45) was not modeled. Second, our structure of the RLC was entirely constructed via homology modeling and the template and target sequences have 43% sequence identity (although the topology of the EF-hand motif is generally conserved across myosin light chains). Finally, there are sparse data for defining the angle that the MyHC tail adopts in between the ELC and RLC binding sites in this chemomechanical state. Our model uses the structure of the scallop isoform under X-ray conditions in the absence of the motor domain. Therefore, results of these simulations may be biased by the initial conformations which may not have placed the RLC at the correct position. Finally, we have simulated a minimal actomyosin complex in solution in the absence of tension. The thin filament is modeled as an actin pentamer that is free to diffuse and tumble: passive axial forces (which would be present in a sarcomere or the laser trap assay) are not present. Similarly, there is no thick filament present: the C-terminal end of MyHC is untethered (which does not correspond to the biological reality of the sarcomere nor to the laser trap. As with any MD simulation the resulting models also suffer from the *sampling problem*: our simulations have probed only a short fraction of time – and there is the potential that larger-amplitude conformational changes due to the A57D mutation may exist but occur with a lower frequency than the simulated timescale.

The two resulting systems were then prepared for molecular dynamics (MD) simulations using the AMBER20 simulation package[10], the ff14SB force field[11], TIP3P water model[12], and the Li and Merz 12-6[13] ion parameters. Missing hydrogen atoms were modeled on using *tleap* and then the systems were placed in periodic boxes (truncated octahedra) and solvated such that the water box extended at least 12 Å beyond any protein atom. Finally, the systems were neutralized with $K^+$ $Cl^-$ counterions and then 120 mM $K^+$ $Cl^-$ ions were added. The systems were then modified using the hydrogen mass repartitioning scheme[14] to enable a 4 femtosecond timestep.

<u>Molecular Dynamics Simulation</u>

Each system was minimized for 10,000 steps divided across three stages in which restraints were placed either on hydrogen atoms, solvent atoms or all backbone heavy atoms ($C_\alpha$, C, N, O atoms). After minimization, systems were heated to 310 K over 300 picoseconds using the canonical NVT (constant number of particles, volume, and temperature) ensemble. During all heating stages, 25 kcal $mol^{-1}$ restraints were present on the backbone heavy atoms ( $C_\alpha$, C, N, O atoms). After the system temperatures reached 310 K, the systems were equilibrated for 5.4 ns over 5 successive stages using the NPT (constant number of particles, pressure, and temperature) ensemble. During equilibration, restraints on backbone atoms were decreased from 25 kcal $mol^{-1}$ during the first stage to 1 kcal $mol^{-1}$ during the fourth stage. During the final equilibration stage, the systems were equilibrated in the absence of restraints. Production dynamics for conventional MD simulations were then performed using the NVT ensemble using a 9 Å nonbonded cutoff and coordinates were saved every picosecond. Single (n=1) 500 nanosecond-long simulations for the WT and mutant AM.ADP_ELC_RLC systems were performed with a 4 femtosecond timestep (enabled due to the repartitioning scheme) (total sampling = 1 microsecond).

Molecular Dynamics Analysis

Unless stated otherwise, analyses were performed on the trajectories subsampled at 10 picosecond intervals. Analysis of the $C_\alpha$ RMSD and $C_\alpha$ RMSF was performed using *cpptraj*[15]. Residue-residue contacts were determined using *cpptraj*: for each timepoint in the simulation, two residues were considered in contact with one another if at least one pair of heavy atoms were within 5 Å of one another. Then we recorded the average percent simulation time each residue pair was in contact for each simulation. To analyze the curvature of the pliant tail, the $C_\alpha$, C, N backbone atom coordinates (residues 773-843 of the 3.AM.ADP simulations; residues were extracted from the simulations and used to fit Bezier curves using the *bezier Python* library[16]. The Bezier curves were defined with a number of points equal to the number of residues included in the tail model, but importantly, each point in the tail corresponds only approximately to each residue number. The curvature was calculated at each pseudo-residue, *i*, on the Bezier curves by fitting a circle to a sliding window of 13 points (13 pseudo-residue positions) centered on pseudo-residue *i*.

Computational Modeling simulating myofilament-level contraction and relaxation.

FiberSim was used to run computational modeling and predict the impact of the A57D MLC1v mutation in human cardiac muscle contraction[17]. The setup file to reproduce the results are added here and accessible in the Github repository (https://doi.org/10.5281/zenodo.18495545).

**Table S1. Differences in interactions made by A57 and D57**

| Interaction Partner | Interaction Partner | Contact Frequency (WT, % time) | Contact Frequency (A57D, % time) |
|---|---|---|---|
| **Generic Residue-Residue Contacts** | | | |
| **57** | MyHC 801 | 98 | 99 |
| **57** | MyHC 804 | 4 | 0 |
| **57** | MyHC 805 | 11 | 13 |
| **57** | MyHC 808 | 24 | 6 |
| **57** | ELC 52 | 12 | 1 |
| **57** | ELC 53 | 100 | 100 |
| **57** | ELC 54 | 100 | 100 |
| **57** | ELC 55 | 100 | 100 |
| **57** | ELC 56 | 100 | 100 |
| **57** | ELC 58 | 100 | 100 |
| **57** | ELC 59 | 100 | 100 |
| **57** | ELC 60 | 100 | 100 |
| **57** | ELC 61 | 99 | 98 |
| **57** | ELC 62 | 9 | 0 |
| **57** | ELC 71 | 4 | 0 |
| **57** | ELC 79 | 13 | 25 |
| **57** | ELC 82 | 1 | 18 |
| **57** | ELC 83 | 87 | 98 |
| **Hydrogen Bonds** | | | |
| RLC:E53@O | RLC:A57@N | 54 | 34 |
| RLC:A57@O | RLC:F61@N | 37 | 22 |
| RLC:A57@O | RLC:L60@N | 22 | 19 |
| RLC:D57@OD1/2 | MyHC:Y801@HH | 0 | 65 |

**Figures.**

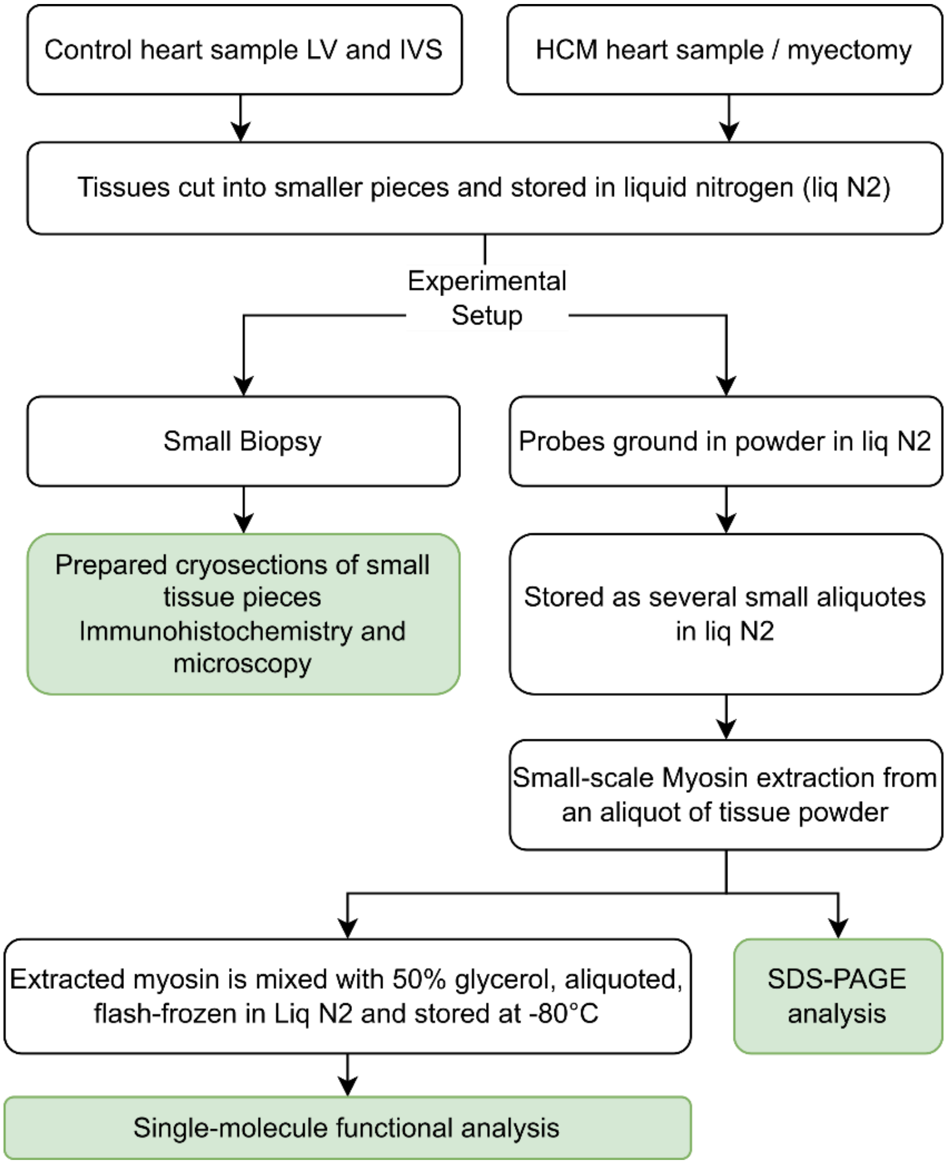


**Figure S1.** Schematic describing workflow adopted for the studies. LV-left ventricle, IVS – intraventricular septum.

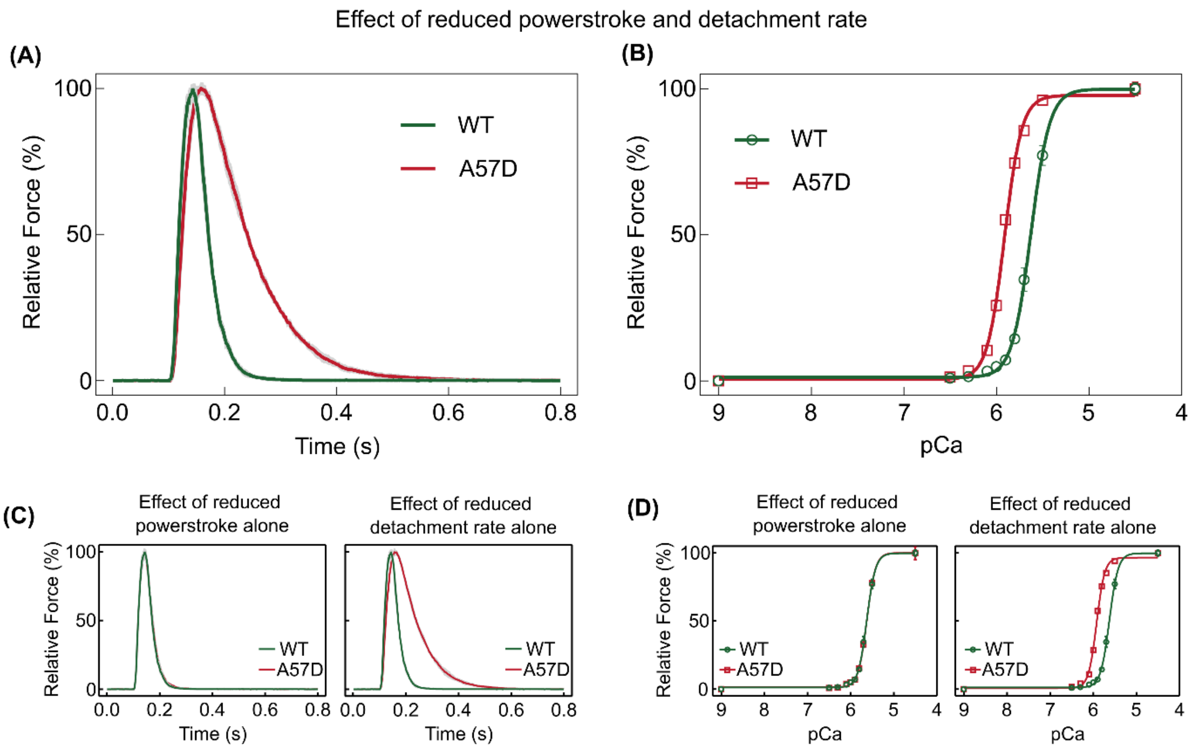


**Figure S2. Simulation of myofilament-level contraction and relaxation parameters: Effect of A57D MLC1v mutation on human myocardial contraction.** Experimentally observed functional alterations in A57D MLC1v myosins at single-molecule level, such as powerstroke size (0.68-fold decrease) and actomyosin detachment rate (0.35-fold decrease), were used to model human cardiac muscle contraction by employing FiberSim software. **(A)** Simulated twitch response to a calcium transient predicts a longer time-to-peak and time for half relaxation of the muscle due to MUT myosins having shorter powerstroke size and reduced detachment rate than WT myosin (n=3, grey lines indicate SD). **(B)** Simulated force-pCa relationship predicts an increase in calcium sensitivity of force generation, with WT having a $pCa_{50}$ = 5.63 and for A57D model had a $pCa_{50}$ = 5.91 (n = 3, error bars indicate SD). **(C)** Simulations for twitch response of muscle with reduced power stroke alone show a relatively similar twitch kinetics between WT and A57D models (left panel). Reduced detachment rate alone produced a slower time-to-peak and half relaxation time in A57D model (right panel). **(D)** Reduced powerstroke size alone did not have a significant effect on calcium sensitivity (left panel). Reduced actomyosin detachment rate alone resulted in a higher $pCa_{50}$ prediction for A57D model (right panel).
